# Oncogene SETDB1’s Dual Role in Endometrial Cancer: Driving Tumor Progression and Immune Escape

**DOI:** 10.1101/2025.04.23.650291

**Authors:** Kiarash Salari, Jiaqing Hao, Ryan Jilek, Matthew Wells, Tianyue Li, Eleanor Johnson, Wendy Meng, Ethan Zibble, Adam Allen, Jack Gilbert, Ryan McLerran, Emma Schuett, Claudia Oliva, Corinne E. Griguer, David K. Meyerholz, Melinda Yates, Henry L. Keen, Xiangbing Meng, Yiqin Xiong, Bing Li, Shujie Yang

## Abstract

Oncogene SETDB1, an H3K9 methyltransferase, drives tumorigenesis in various cancers. Using endometrial cancer (EC) as a model, we discovered SETDB1’s dual mechanisms in driving EC tumorigenesis and mediating immune evasion. SETDB1 knockout (SETDB1^-/-^) tumor-bearing mice exhibited prolonged survival of up to 100 days. Transcriptomic profiling of SETDB1^-/-^ EC cells revealed decreased expression of oncogenes (*POLR2A, MSH6, FLNA*) and increased expression of tumor suppressor genes (*PGR, RERG, ZNF582*), which indicates that SETDB1 intrinsically promotes EC tumor proliferation by regulating these downstream genes. SETDB1 repressed repeat elements and the interferon pathway, mediating immune evasion extrinsically by inhibiting anti-tumor macrophage infiltration. ChIP-seq analysis showed SETDB1 binding at pericentromeric regions on many chromosomes and numerous ZNFs. Loss of SETDB1 resulted in abnormal cell division. SETDB1^-/-^ tumors displayed reduced proliferation markers (Ki67, pHH3) and increased macrophage infiltration.

Mechanistically, SETDB1 promotes CD47 (a don’t-eat-me signal) and represses CCL5 and CXCL9 (macrophage and T-cell recruiting chemokines), contributing to immune evasion. M1-like macrophages killed more SETDB1^-/-^ cells in co-culture. Additionally, SETDB1 knockout in mouse EC cells reduced tumor growth in C57BL/6 mice, with increased macrophage and CD4+ T-cell infiltration. Our results indicates that elevated SETDB1 and its target genes can predict higher tumor grade and worse survival, suggesting that targeting SETDB1 could be a promising therapeutic strategy for EC.

## Introduction

Oncogene SETDB1 (SET Domain Bifurcated Histone Methyltransferase 1, ESET, KMT1E) is a histone H3K9 methyltransferase responsible for mono-, di-, and tri-methylation of histone H3K9^1, 2^. The SET domain—originally identified in **<u>S</u>**u(var)3-9, **<u>E</u>**nhancer of zeste [E(z)], and **<u>t</u>**rithorax (trx)—is crucial for H3K9 trimethlylation^3^. Overexpression of SETDB1 has been reported in many cancer types^4, 5^, and is linked to worse patient outcomes due to elevated SETDB1 mRNA expression from high copy number^4, 6^,. SETDB1 is located on the chromosome 1q21.3. Chromosome 1q amplification, a known hotspot, has been reported to predict worse prognosis in patients across various cancer types^7, 8, 9, 10^. Amplified SETDB1 promotes heterochromatin formation and silencing tumor suppressors^11^. We and other groups reported that SETDB1 executes five critical functions intrinsically: (1) epigenetic effects, (2) PML-NB (promyelocytic leukemia nuclear body) formation, (3) retroelement silencing, (4) inhibition of cell cycle arrest, and (5) cell proliferation^4^. Furthermore, SETDB1 also plays a role in immune suppression and tumor immunogenicity extrinsically as summarized by our group and others^5, 12, 13^. Specifically, it (1) alters lymphocyte and cytokine expression by suppressing the production and infiltration of antitumor immune cells, (2) facilitates tumor immune escape, (3) disrupts the type I interferon (IFN) pathway, and (4) decreases ICB (immune checkpoint blockade) effectiveness. Despite advances, there remains a substantial gap in understanding how SETDB1 contributes to tumor progression via these two mechanisms, specifically, whether SETDB1 will employ both mechanisms in a single tumor system.

Our group focuses on endometrial cancer (EC), the most common gynecologic cancer in the US and globally, originating from the inner lining of the uterus. EC causes approximately 91,640 deaths annually worldwide^14^. Unlike other cancers, EC is the only type with declining survival rates over the past 40 years^15^, highlighting the urgent need for improved treatment. Two studies on EC, proteogenomic and Pan-Cancer data analyses, have revealed that SETDB1 is overexpressed ^16, 17^. However, neither analysis focuses on SETDB1’s oncogenic mechanisms. Given the critical unmet need to improve clinical outcomes for EC patients, defining SETDB1’s oncogenic mechanisms in EC represents an important advancement. Using EC as a model, we aimed to identify the molecular mechanisms driven by SETDB1 in EC by utilizing established EC cell lines, xenograft and syngeneic models, and patient samples.

To understand SETDB1’s role in EC tumorigenesis, we knocked out SETDB1 in EC cell lines and grew tumors in both immunocompromised and immunocompetent mice. Depletion of SETDB1 greatly slowed cell proliferation and tumor formation, and increased macrophage infiltration in SETDB1^-/-^ tumors. Potential mechanisms of SETDB1-mediated tumorigenesis include: (1) an intrinsic mechanism: promoting oncogene transcription and silencing tumor suppressors, binding to pericentromeric regions of chromosomes, facilitating proper cell division; and (2) an extrinsic mechanism: establishing an immune-suppressed state, promoting the “don’t-eat-me” signal CD47, blocking macrophages infiltration, inhibiting repeat elements expressions and IFN pathways, and repressing macrophage and T-cell chemokines CCL5 and CXCL9. Our data demonstrates that SETDB1 drives EC tumorigenesis and fosters immune evasion.

## Results

### Overexpressed SETDB1 predicts worse survival and active gene transcription signature

Elevated SETDB1 expression has been reported across many cancer types including EC^3, 4^. Using the Pan-Cancer TCGA (The Cancer Genome Atlas) and CPTAC (Clinical Proteomic Tumor Analysis Consortium) dataset, elevated SETDB1 was found in most tumor types at both mRNA and protein levels (Supplementary Fig. 1a-b). In the CPTAC dataset, SETDB1 exhibited the highest expression in EC among the eight tested tumor types (Supplementary Fig. 1c). To verify SETDB1 overexpression in EC, we used two available datasets: EC-TCGA TARGET and GDC TCGA. First, in EC-TCGA GTEX dataset, we confirmed substantially higher SETDB1 expression in tumor samples (n=57) (mean =11.31) compared to normal tissues (n=78) (mean = 10.97) (p < 0.0001) (Fig. 1a). In the GDC TCGA dataset, we confirmed significantly higher SETDB1 expression in tumors (n=589) than normal (n=35) (p=0.0139) (Fig. 1b).

**Fig. 1.**
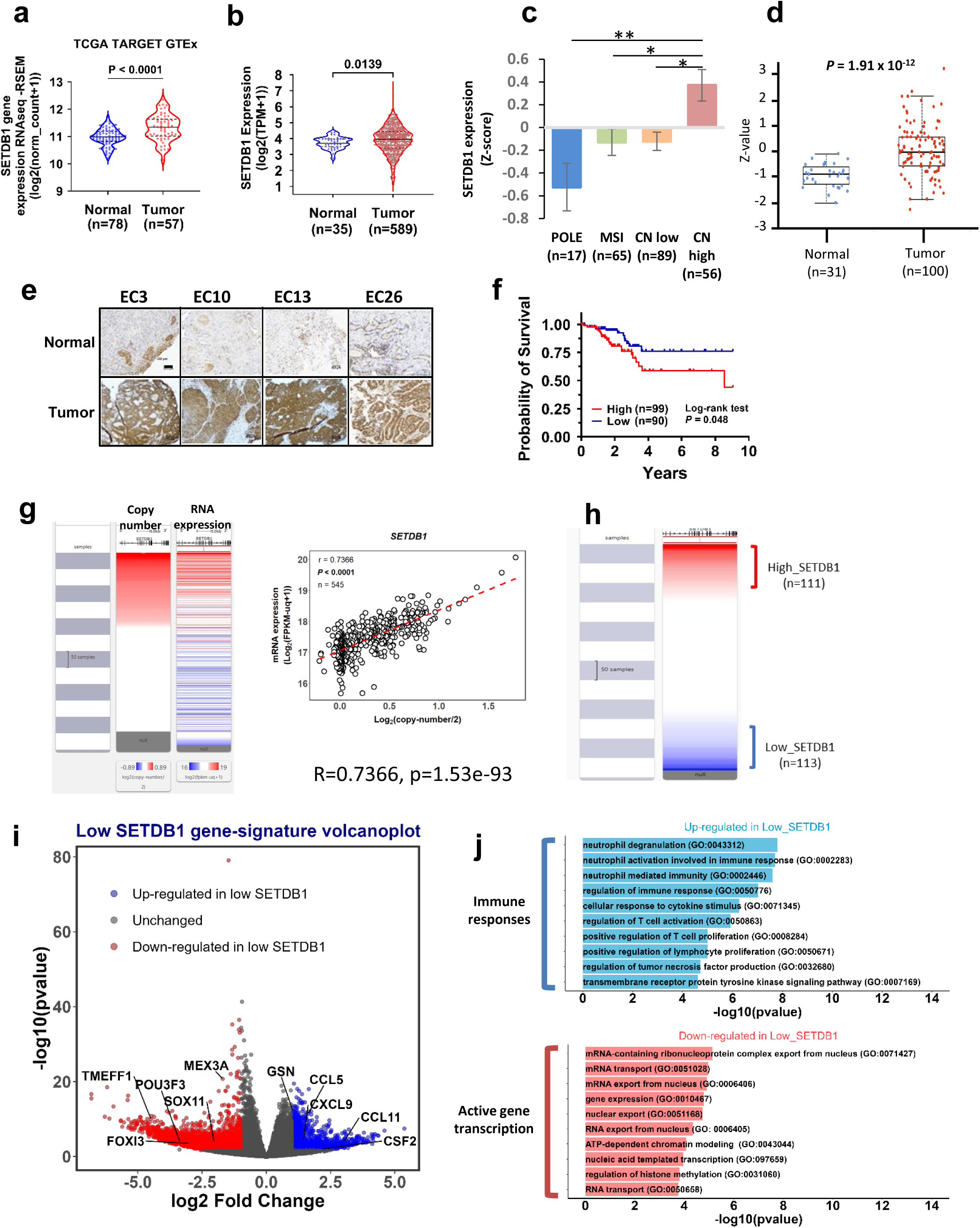
Overexpressed SETDB1 predicts worse survival and active gene transcription signature. a, b, **d** SETDB1 expression comparison at mRNA and protein level in normal and tumor tissues across different datasets: EC-TCGA TARGET mRNA expression (normal, n=78; Tumor, n=57) (**a**) EC GDC TCGA mRNA expression (normal, n=35; Tumor, n=589) (**b**) CPTAC protein expression (normal, n=31; Tumor, n=100) (**d**). The solid middle line in violin plots represents median and the dashed lines are quartiles. Boxplots show the distribution of data across groups. The central line represents the median value. The low and high end of the box represent 25th and 75th percentiles respectively. **c** SETDB1 mRNA expression across different UCEC subtypes of POLE (n=17), MSI (n=65), CN low (n=89) and high (n=56) groups. **e** Representative tissue IHC staining of SETDB1 comparing normal adjacent and tumor tissues. Scale bar represents 100μm. **f** Kaplan Meier survival curve for low (n=90) and high (n=99) SETDB1 expressing patients. **g** Pearson’s correlation between SETDB1 DNA copy number and its RNA expression utilizing TCGA dataset (n=545). **h** Heatmap displaying separation between high (n=111) and low (n=113) SETDB1 groups of patients’ samples. **i** Volcano plot for EC TCGA differential expression analysis on downregulated (red) and upregulated (blue) genes in Low SETDB1 group of patients. **j** Gene Ontology analysis on up (blue) and downregulated (red) genes in low SETDB1 groups. Statistical tests: unpaired students’ *t* test (**a, b**, **c, d**), log-rank test (**f**), Pearson’s correlation coefficient (**g**), Wald test (**i**), Fisher’s exact test (**j**), **c** *, **, *P* < 0.05, *P* < 0.01.

The TCGA classifies EC into four molecular subtypes (1) PolE/ultra-mutated (<u>Pol</u>ymerase <u>E</u>psilon mutation), (2) MSI/ hypermutated (<u>M</u>icro<u>s</u>atellite Instability), (3) CNL/NSMP (<u>C</u>opy <u>N</u>umber <u>L</u>ow/<u>N</u>o <u>S</u>pecific <u>M</u>olecular <u>P</u>rofile), and (4) CNH/p53abn (<u>C</u>opy <u>N</u>umber <u>H</u>igh and <u>p53 abn</u>ormal) ^18, 19^. We found that SETDB1 is highest in the CNH group (Fig. 1c) which is the most aggressive and poorly immunogenic subtype of EC. Elevated SETDB1 expression at mRNA levels (Fig. 1a-c) were further confirmed at protein levels in tumors using the CPTAC dataset (Fig. 1d).

To visualize SETDB1 expression using immunohistochemistry (IHC) staining, 76 patient tumors with matched adjacent non-malignant tissues were collected from the University of Iowa Hospital and Clinics, confirming higher SETDB1 protein expression in many tumors (Fig. 1e). Using the EC-TCGA dataset, we verified that patients with high SETDB1 expression had more unfavorable outcomes (p=0.048) (Fig. 1f, Supplementary Fig. 1d). Additionally, we confirmed a strong positive correlation (r=0.7366, p=1.53e-93) between copy number and mRNA expression (Fig. 1g) in EC, consistently observed in other cancers^7^.

To understand the gene expression landscape of high and low SETDB1 groups, we conducted transcriptomic analysis using the EC-TCGA. We divided EC-TCGA patients into high (n=111) and low (n=113) SETDB1 expressing subgroups and conducted differential gene expression analyses (Fig. 1h). Patients with high SETDB1 levels presented greater expression of genes related to active transcription signature, including many mRNA transcription regulators (Fig. 1i, j). In contrast, those with lower SETDB1 levels expressed high levels of immune response genes, such as CCL5, CXCL9, and CCL11 (Fig. 1i, j). These findings highlight the critical functions of SETDB1 in EC and provide insights into the potential downstream genes and pathways influenced by SETDB1 expression levels in EC.

### SETDB1 depletion decreases EC cell proliferation *in vitro*

To uncover the pivotal role of SETDB1 in driving EC cell proliferation, we started our investigation with three established EC cell lines and expanded to our three novel primary patient-derived EC cell lines (PDCs). We designed three sgRNA to knockout SETDB1 using CRISPR-Cas9 in Ishikawa (poorly differentiated), ECC1 (well-differentiated), and Hec50 (serous) cell lines. We successfully generated numerous knockout clones, achieving near-total depletion of the SETDB1 protein (Supplementary Fig. 2a, c, e).

Our initial *in vitro* experiments revealed the knockout cells exhibited a markedly reduced growth rate compared to wildtype (WT, non-target sgRNA) cells, though with some variability (Supplementary Fig. 2b, d, f). We selected one knockout clone per sgRNA (totaling three knockout clones) and confirmed SETDB1 depletion (Supplementary 3a, c, e).

To delve deeper, we monitored the proliferation of these EC cells over four days (Supplementary Fig. 3b, d, f). Remarkably, SETDB1 depletion consistently inhibited cell proliferation across all tested knockout clones. Specifically, Ishikawa SETDB1^-/-^ clones showed approximately 10-30% reduction compared to WT (Supplementary Fig. 3b). Similarly, ECC1 and Hec50 cells with SETDB1 depletion demonstrated a dramatic 30-60% decrease in proliferation across all tested clones (Supplementary Fig. 3d, f). Our results repeatedly demonstrated that SETDB1 is crucial in driving EC cell proliferation in three different EC subtypes (Supplementary Fig. 1, 2b, d and f).

To extend these findings to patient tumors, we utilized PDCs to investigate SETDB1’s role in EC cell proliferation. We selected PDC4 (Grade 2 EC), PDC8 (serous EC), and PDC10 (recurrent EC) as representative examples. We confirmed successful SETDB1 knockout, which led to a significant reduction in cell proliferation by 30-40% (Supplementary Fig. 3h-j). This finding confirmed the critical function of SETDB1 in promoting EC cell proliferation in patient models.

In summary, our data from SETDB1 depletion in six EC cell models underscore its critical role in driving EC cell proliferation.

### Depleting SETDB1 significantly slows down tumor growth and prolongs mice survival *in vivo*

To investigate whether oncogene SETDB1 promotes tumor growth, we injected SETDB1^-/-^ EC cells (Ishikawa, ECC1, and Hec50) into NSG mice (NOD.Cg-Prkdcscid Il2rgtm1Wjl/SzJ) subcutaneously on their left and right flanks. SETDB1 depletion was confirmed by western blotting (as shown in Fig. 2a, d, g). Our results showed that SETDB1 knockout significantly slowed tumor growth rate across all cell lines (Fig. 2b, e, h). The most significant reduction was observed in the Ishikawa cell line. Remarkably, mice bearing SETDB1^-/-^ tumors exhibited a substantially extended survival, up to 100 days for Ishikawa and ECC1, and 60 days for Hec50 (Fig. 2c, f, i).

**Fig. 2.**
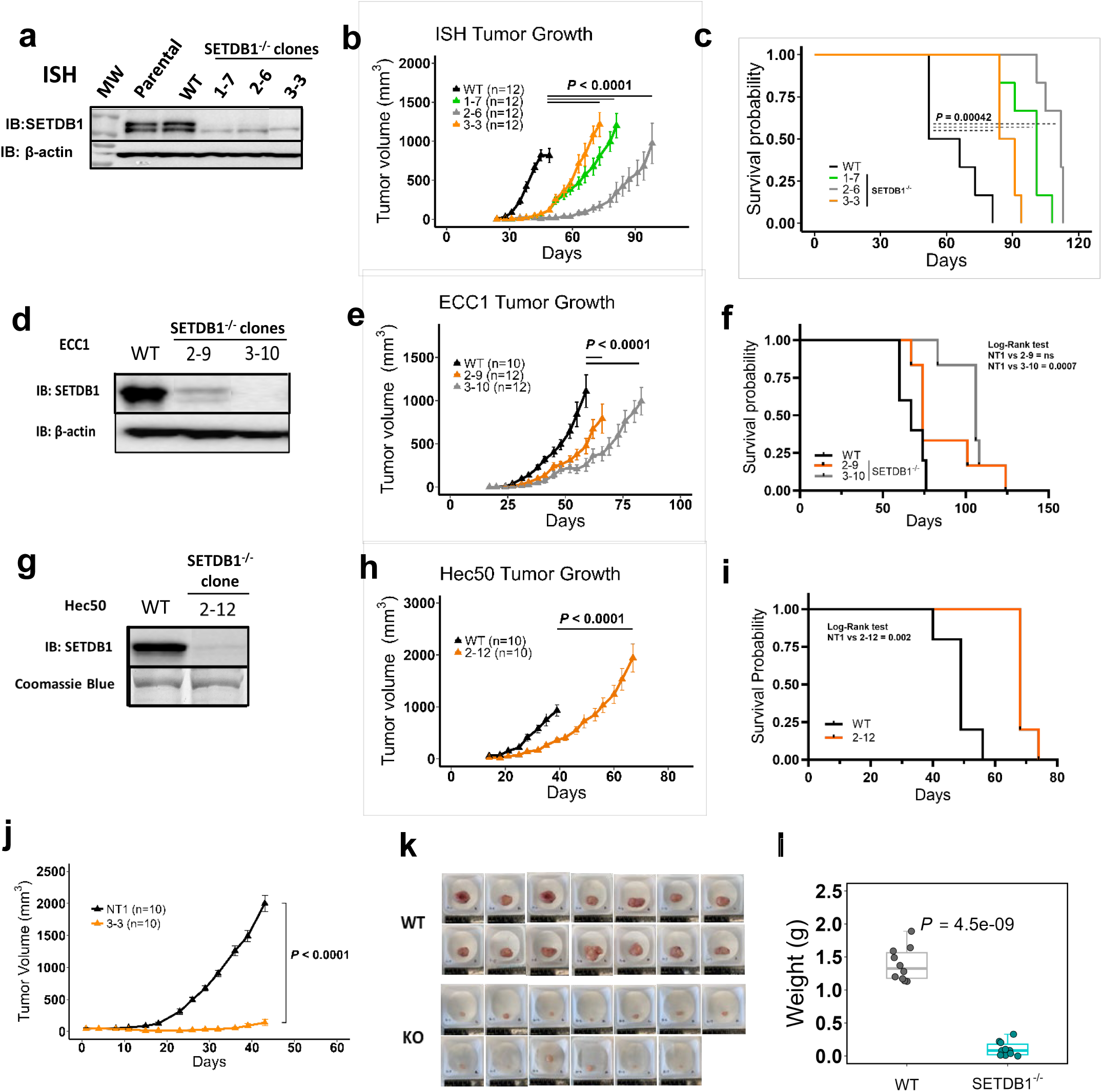
knockout SETDB1 decreases tumor growth significantly *in vivo* and prolongs NSG mice survival. a,d,g. SETDB1 immunoblot for SETDB1 knockout clones and wildtype for Ishikawa (ISH), ECC1, and Hec50 cell lines respectively. **b,e,h** Mean tumor volume measurements over time for SETDB1 knockout clones and wildtype subcutaneously injected in female NSG mice for cancer cells: ISH (n=12/group for WT, 1-7, 2-6, and 3-3) (**b**), ECC1 (n=10/group for WT, n=12/group for 2-9, 3-10) (**e**), Hec50 (n=10/group for WT, 2-12) (**h**) with n representing number of total injection sites and/or tumors. **c,f,i** Kaplan Meier survival curves for NSG mice injected with SETDB1 knockout clones and WT for cancer cells: ISH (n=6/group for WT, 1-7, 2-6, and 3-3) (**c**) ECC1 (n=5/group for WT and n=6/group for 2-9, 3-10) (**f**) Hec50 (n=5/group for WT, 2-12) (**i**) with n representing number of mice in each group. **j** Mean tumor volume measurements over time for ISH wildtype and 3-3 knockout clone with experiment ending when wildtype tumor reaches endpoint (n=10/group for WT, 3-3) with n representing number of total tumors. **k** Tumor pictures showing relative tumor sizes of wildtype and SETDB1 knockout clone 3-3 from experiment in **j**. **l** Tumor weights measurements of wildtype and SETDB1^-/-^ tumors in **k**. Boxplot showing the distribution of data on the plot and extended to the highest and lowest limits of the datapoints. Middle horizontal line on the boxplot represents the median. **b-h, j,** Data shown as mean ± SEM. Statistical tests: A non-linear fit (Exponential growth model as log(population)) to the curves was generated and the fitted curves were compared using extra sum-of-squares F test to calculate the p value (**b, e, h**), Log-Rank test (**c, f, i**) Student’s t-test (**j**) Welsh’s two sample t-test (**l**).

To further confirm these findings, we conducted a fixed time point experiment for Ishikawa cells. The SETDB1^-/-^ tumors demonstrated significantly smaller volumes (93.1% reduction) and weights (92.2% decrease) (Fig. 2j, k, l). These experiments were performed multiple times with consistent and reproducible results (Supplementary Fig. 4a, b). Altogether, our data illustrates that knockout of SETDB1significantly inhibits tumor growth and prolongs survival in NSG mice.

### SETDB1 enhances oncogenes and represses tumor suppressor genes

To determine downstream genes controlled by SETDB1, we conducted transcriptomic analysis on our three Ishikawa knockout clones and WT control (shown in Fig. 2a, b). We compared differential gene expression and observed 677 shared genes significantly altered in all SETDB1^-/-^ clones (Fig. 3a). Gene set enrichment analysis (GSEA) revealed that genes associated with Mitotic Spindle and G2M Checkpoint were the most significantly downregulated biological processes in SETDB1^-/-^ cells (Fig. 3b). Conversely, some genes belonging to IFNα and IFNγ pathways were among those upregulated in SETDB1^-/-^ (Fig. 3c). RNA-sequencing revealed two groups of representative genes altered in SETDB1^-/-^ cells: Group 1, containing upregulated genes (repressed by SETDB1) mainly functioning as tumor suppressors; and Group 2, containing downregulated genes (promoted by SETDB1) that are mainly oncogenes involved in active gene transcription and cell proliferation (Fig. 3d, e).

**Fig. 3.**
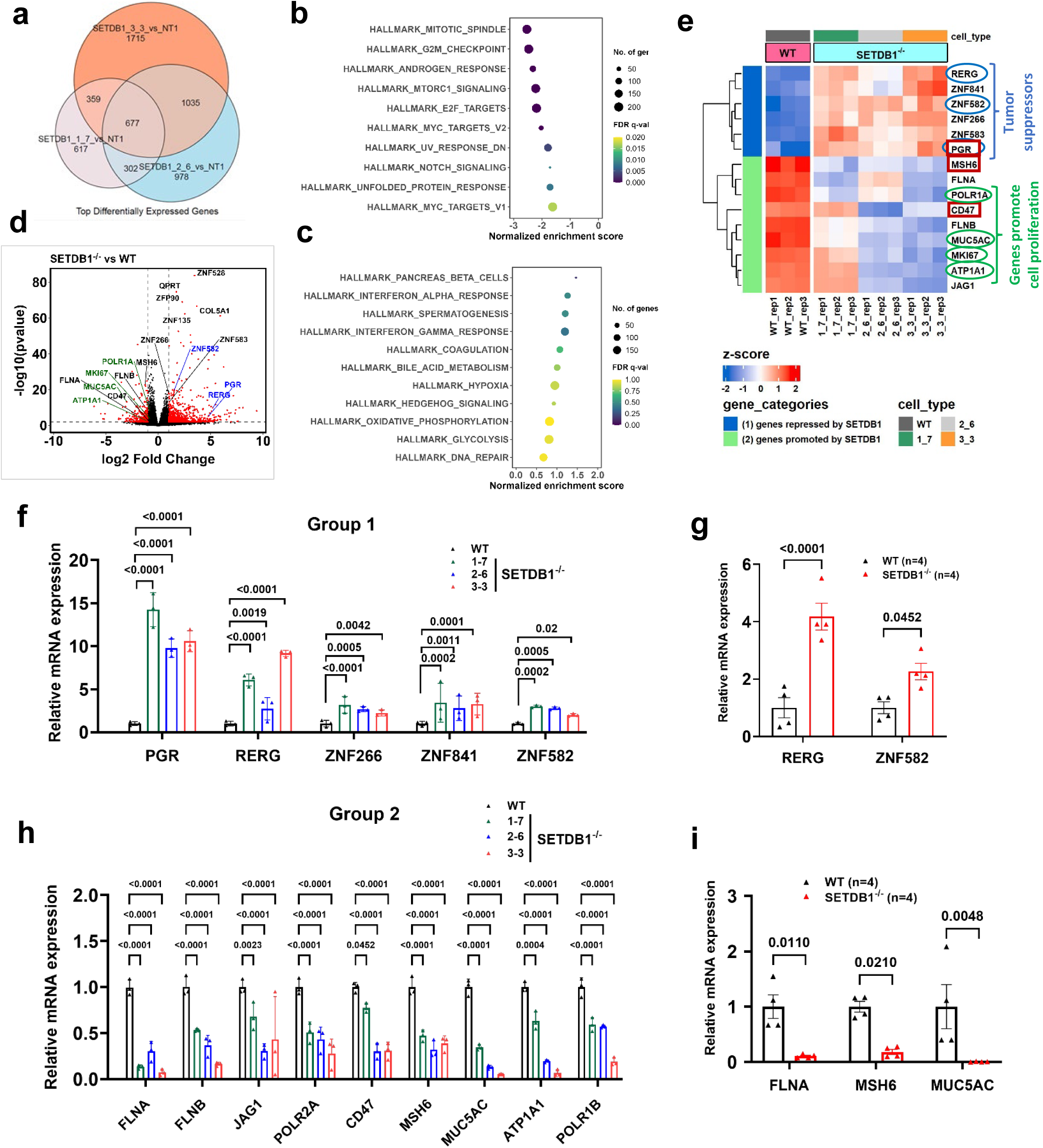
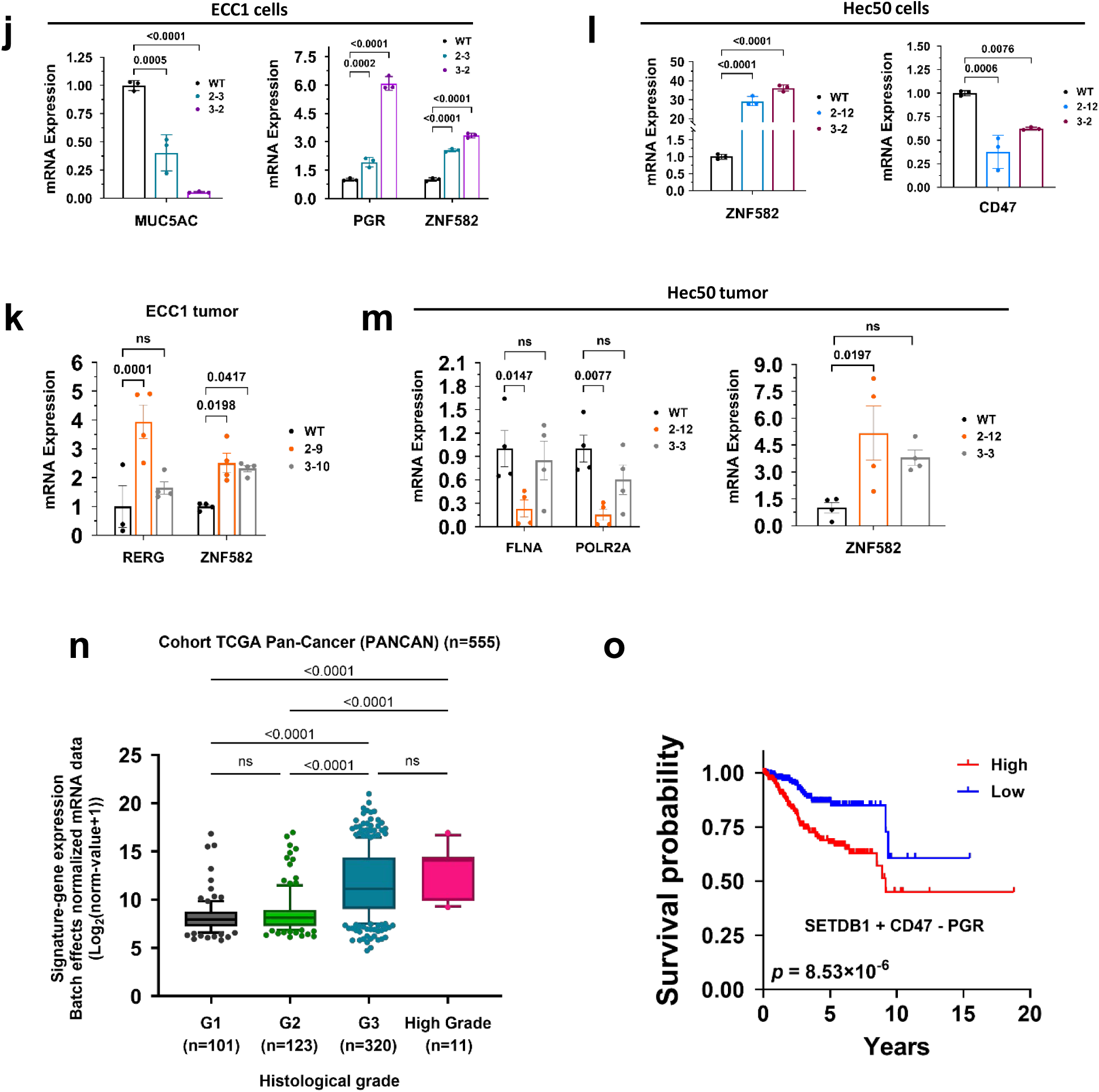
SETDB1 promotes oncogene expression and inhibits tumor suppressor gene expression (RNA-seq) **a** Venn diagram demonstrating gene expression changes for different SETDB1 knockout clones and their overlaps with one another. **b, c** Gene Set Enrichment Analysis (GSEA) depicting biological processes downregulated or upregulated respectively in SETDB1^-/-^ clones compared to wildtype Ishikawa cells. **d** Volcano plot for differentially expressed genes in all ISH SETDB1 knockout clones compared to wildtype; red represent genes with p_adj_<0.1 and abs(log2FoldChange)>1. Green labeled and blue labeled genes are known oncogenes and tumor suppressors respectively. **e** Heatmap presenting gene expression changes (presented as z-score) of different SETDB1 knockout clones (n=3 biological replicates). The blue circles indicate tumor suppressors, and the green circles highlight genes promoting cell proliferation. Red boxes represent genes with SETDB1 activity signatures. **f, h** qPCR verification for differentially expressed Group 1, and 2 genes in SETDB1^-/-^ clones respectively (n=3 biological replicates). **g, i** qPCR analysis of selected Group 1 and 2 genes in wildtype and SETDB1^-/-^ tumors respectively (n=4 biological replicates). **j, l** verified qPCR results on Group 1 and 2 genes in SETDB1 knockout and wildtype ECC1 and Hec50 cell lines respectively (n=3 technical replicates). **k, m** verified qPCR results on Group 1 and 2 genes in SETDB1 knockout and wildtype ECC1 and Hec50 tumors respectively (n=4 biological replicates). **n** SETDB1 and its downstream genes (MSH6 and PGR) mRNA expressions differentiating EC tumor grades in patients. **o** Kaplan Meier survival curve characterizing patients based on SETDB1 and its downstream genes (CD47, and PGR) mRNA expressions predicting EC patients’ outcome. For **n**, boxplots are expanded from 10th to 90th percentiles. The low and high end of the box represent 25th and 75th percentiles respectively. Middle horizontal line represents the median. **f-m**, Data shown as mean ± SD. Statistical tests: Wald test (**d**), Two-way ANOVA with post hoc Dunnett’s multiple comparisons test (**f, h, j (2^nd^ part), k, m (1^st^ part)**), One-way ANOVA with post hoc Dunnet’s multiple comparison test (**j (1^st^ part), l, m (2^nd^ part)**), Two-way ANOVA with Sidak’s multiple comparisons test (**g, i**), Kruskal-Wallis test (**n**), Log-Rank test (**o**).

Representative Group 1 genes repressed by SETDB1 include *PGR* (Progesterone Receptor), *RERG* (RAS-Like Estrogen-Regulated Growth Inhibitor), and *ZNF582* (Zinc finger protein 582). SETDB1 depletion led to the upregulation of these genes. Among the most upregulated Group 1 genes were *PGR* and *RERG,* demonstrating a remarkable 10-14 fold and 3-9 fold upregulation, respectively (Fig. 3f).

*PGR* is reported as an ultimate tumor suppressor gene in EC^20^. *RERG* is a tumor suppressor in breast^21^, prostate^22^, nasopharyngeal^23^, and glioma^24^ cancers. *ZNF582,* a zinc-finger transcription factor, is one of the few reported tumor suppressors among zinc finger proteins^25^. Next, we used the SETDB1^-/-^ tumors to verify the expression of these target genes and recapitulated SETDB1-mediated *RERG* and *ZNF582* repression (Fig. 3g). To generalize the SETDB1 regulated downstream genes, we used additional SETDB1^-/-^ clones and verified upregulation of *RERG* and *ZNF582* by 40-60 fold and 4-5 fold, respectively (Supplementary Fig. 5a).

Next, we evaluated the clinical relevance of SETDB1 target Group 1 genes in EC. These genes showed higher expression in normal tissues in TCGA (Supplementary Fig. 5b, c) and CPTAC (Supplementary Fig. 5d) datasets. Using the TCGA dataset, Group 1 genes correlated with better patient survival overall (Supplementary Fig. 6a).

In contrast, SETDB1 promoted Group 2 genes, including *FLNA* (Filamin A), *POLR2A* (RNA polymerase II), *CD47*, *MSH6* (mutS homolog 6), and *MUC5AC*. SETDB1 depletion resulted in downregulation of these genes. *FLNA*, *POLR2A,* and *MUC5AC* were among the most downregulated genes, with fold changes ranging from 0.07-0.29, 0.24-0.5, and 0.05-0.35, respectively (Fig. 3h). *FLNA* is crucial for maintaining cell shape and movement^26^. *POLR2A* is an essential transcription gene controlling rapid cell proliferation^27^. *CD47*, a “do-not-eat me” signal, is often upregulated in cancer to prevent immune clearance^28^. *MSH6*, a critical mismatch repair gene, is associated with worse survival in many cancers despite its mutation increasing cancer susceptibility^29^. *MUC5AC* promotes cancer progression and metastasis in various cancers^30^. Consistently, we confirmed the downregulation of *FLNA, MSH6,* and *MUC5AC* in SETDB1*^-/-^* tumors (Fig. 3i). To generalize the SETDB1 regulated downstream genes, we used additional SETDB1^-/-^ Ishikawa clones and verified *POLR2A* downregulation by 0.25-0.75 fold in clone sg1-12 and sg3-12 (Supplementary Fig. 5a).

Next, we evaluated the clinical relevance of SETDB1 target Group 2 genes in EC. These genes showed lower expression in normal tissues in TCGA (Supplementary Fig. 5e, f) and CPTAC (Supplementary Fig. 5g) datasets. Using the TCGA dataset, Group 2 genes correlated with unfavorable patient survival overall (Supplementary Fig. 6b).

To further validate our findings, we examined two additional cell lines, ECC1 and Hec50. In SETDB1^-/-^ ECC1 cells, *MUC5AC* was downregulated, while *PGR* and *ZNF582* were upregulated (Fig. 3j). In SETDB1^-/-^ Hec50 cells, *ZNF582* was upregulated, and *CD47* was downregulated (Fig. 3l). Furthermore, we recapitulated similar results in SETDB1^-/-^ tumors. With *RERG* and *ZNF582* upregulated in ECC1 tumors (Fig. 3k), while *ZNF582* was upregulated, and *FLNA* and *POLR2A* were downregulated in SETDB1^-/-^ Hec50 tumors (Fig. 3m). These findings demonstrate that while SETDB1 regulates a range of target genes, *ZNF582* regulation is consistently observed across multiple cell types, underscoring its potential as a key downstream effector of SETDB1. Other target genes exhibit cell type specific regulation, reflecting the contextual roles of SETDB1.

To find the clinical application of SETDB1 and its target genes, we identified four downstream genes, along with SETDB1, providing a predictive model for EC tumor grades and patient outcomes. We named these genes “SETDB1 activity signature” genes. The three-gene signature “SETDB1+MSH6-PGR” can distinguish EC grades from one another (Fig. 3n). The three-gene signature, “SETDB1+ CD47-PGR” correlates with worse patient prognosis (p=3.016e-7) (Fig. 3o). These data emphasize the importance of SETDB1 in EC progression and outcomes.

### SETDB1 binds Scaffold Attachment Regions (SARs) at centromeres and represses ZNF genes via H3K9me3 deposition at their promoters

SETDB1 has been reported to regulate gene expression by directly binding to the gene’s promoter region in different systems^2, 31, 32^. To explore SETDB1 binding sites on a whole genome level, we conducted ChIP-seq for H3K9me3 and SETDB1 on Ishikawa cells. ChIP-seq data surprisingly revealed that SETDB1 binds dominantly to the SARs of pericentromeres on chromosome (chr) 3 (Fig. 4a) and chr 4 (Fig. 4b) with a striking high peak, 90,000 (chr 3) and 2,000 (chr 4). Knockout of SETDB1 abolishes its binding, which we termed “irreplaceable constant binding.” A 2022 Science paper identifies SAR, termed “Hsat1A,” on chr 3, 4, 8, 13, 14, 21 and 22^33^. In addition to chr 3 and 4, we confirmed the prominent binding peak of H3K9Me3 and SETDB1 on chr 8, 14, and 22, with less significant peaks on chr 13 and 21 (Fig. 4 c-e). While H3K9me3 has been reported to bind centromeres^34^, we found that SETDB1 binding to centromeres in human cancer cells, highlighting a novel and unexpected role for SETDB1.

**Fig 4.**
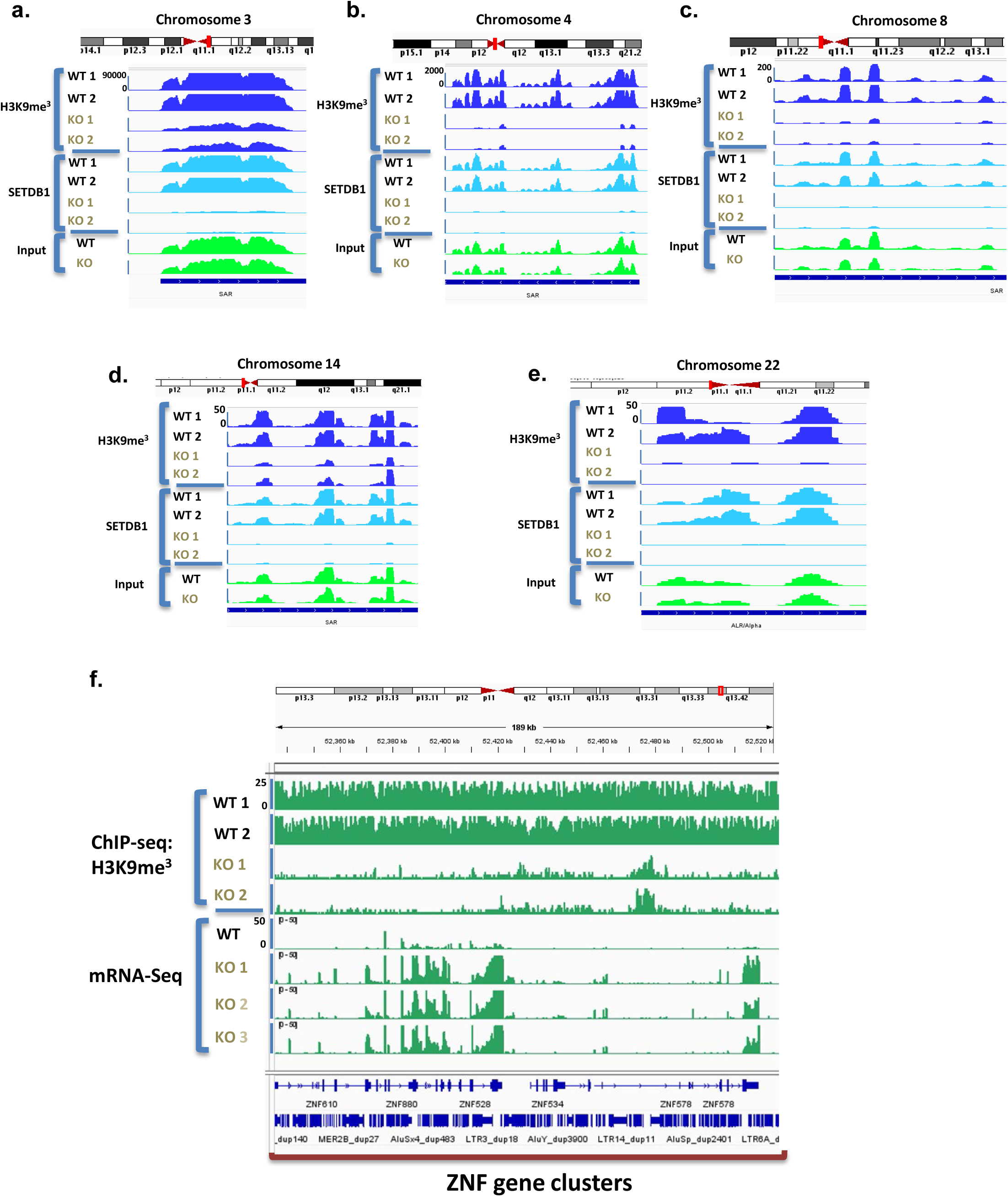
SETDB1 directly binds SARs on the centromeres and deposits H3K9me3 on ZNF genes promoters. a-e. ChIP-seq H3K9me3 and SETDB1 peaks related to enrichment for genomic DNA sequences corresponding to different peri-centromeric regions within chromosomes 3, 4, 8, 14, and 22 respectively for both wildtype and SETDB1 knock cells. The Y-axis is labeled only on the first sample, but the scale is consistent across all samples. **f** ChIP-seq H3K9me3 and RNA-seq peaks on an enriched ZNF gene cluster region within chromosome 19 for both wildtype and SETDB1 knockout cancer cells.

Besides SETDB1 binding to the pericentromere, our ChIP-seq illustrated that SETDB1-mediated H3K9Me3 directly binds to many ZNFs to repress their expression; SETDB1 depletion leads to ZNF upregulation (Fig. 4f). Interestingly, SETDB1 deposits the H3K9Me3 mark on ZNFs but then dissociates from the site (Supplementary Fig. 7a). To verify H3K9Me3 binding to ZNF, ChIP-PCR was conducted. We found that H3K9Me3 binds directly to ZNF266, ZNF841, ZNF582, while this binding was lost in the SETDB1^-/-^ cells (Supplementary Fig. 7b). To validate this result, we repeated this experiment with one additional clone and achieved similar results (Supplementary Fig. 7c). We termed this binding pattern as “irreplaceable transient binding,” highlighting SETDB1’s role in depositing repressive marks and then leaving, ensuring gene repression is maintained.

### Knockout of SETDB1 causes abnormal chromosome division during mitosis

While we know SETDB1 binds to the SAR of the pericentromere, we lack alternative verification methods. We were inspired by two papers on Setdb1 depletion in mouse germlines that reported that loss of Setdb1 impairs meiosis and mitosis^35^. In mouse oocytes and early embryos, it was reported that loss of Setdb1 significantly impaired kinetochore-spindle interactions, bipolar spindle organization, and chromosome segregation. Thus, we applied immunofluorescent (IF) labeling using β-tubulin for the tubulin spindle, γ-tubulin for the centrosome, and DAPI for the chromosome. In contrast to the Ishikawa wild type cells (Fig. 5a, a1-3, Supplementary Fig. 8a, a1-4), we were excited to find that SETDB1^-/-^ cells showed multiple γ-tubulin poles during mitosis, generating three, four, and even six poles of γ-tubulin (Fig. 5b, b1-6, Supplementary Fig. 8b, b1-6). These multiple poles led to either micronuclei (Supplementary Fig. 8b, b7) or three poles of dividing cells (Fig. 5b, b2, Supplementary Fig. 8b, b8-9). Overall, SETDB1^-/-^ generated 12% of abnormal mitotic cells, while the wild type had only 2.5% of abnormal mitotic cells (Fig. 5c).

**Fig 5.**
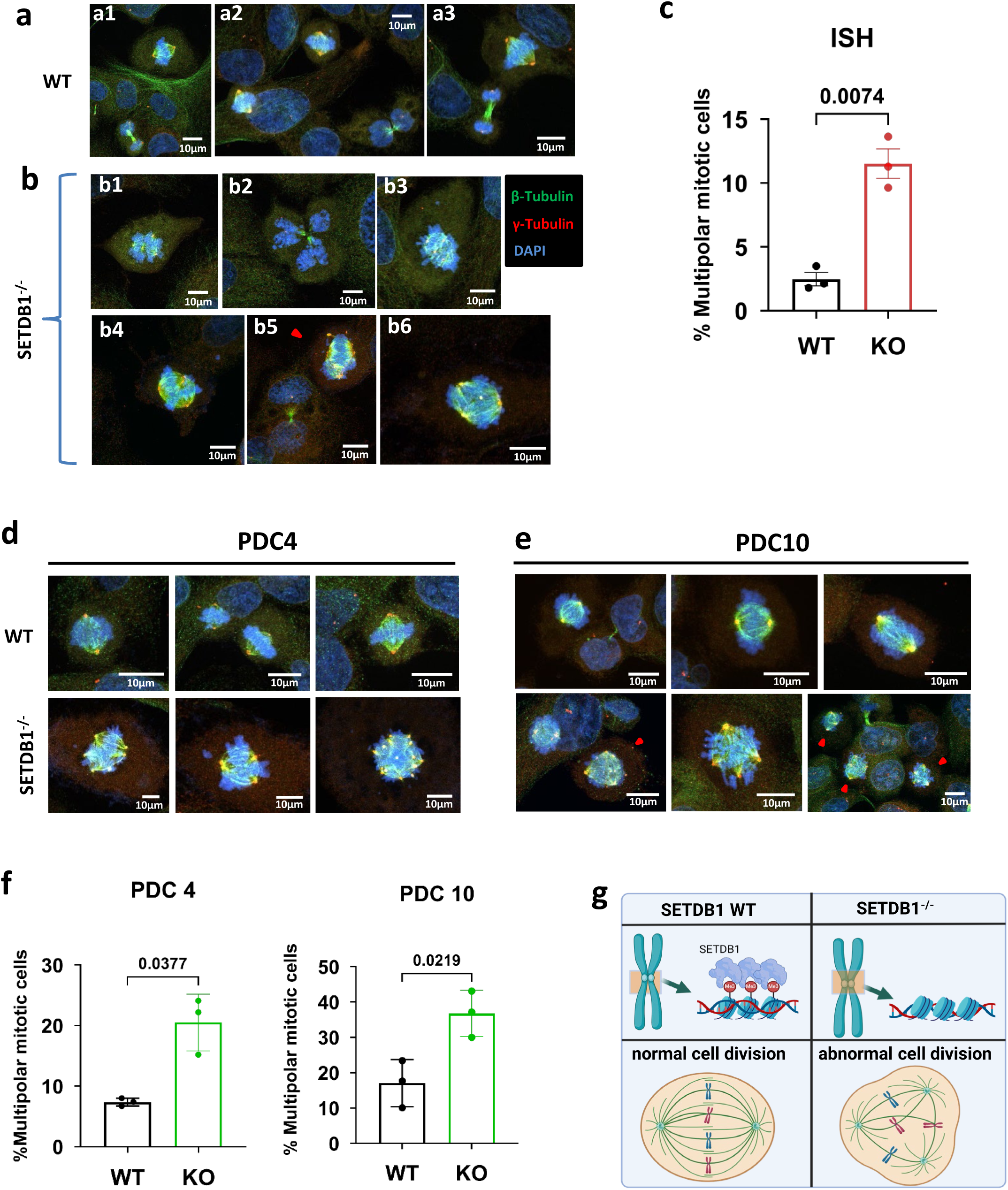
Knockout SETDB1 leads to defect in chromosome segregation. a, **b** Representative Immunofluorescence staining images of bipolar and multipolar mitotic cells in wildtype and SETDB1 knockout Ishikawa cells respectively. Red: γ-tubulin, green: β-tubulin, blue: genomic DNA. Red arrowhead points to the abnormal mitotic cells. **c** Percent multipolar mitotic cells quantification in wildtype and SETDB1 knockout cells (n=3 biological replicates). More than 80 mitotic cells in different fields were analyzed for abnormalities in each biological replicates within wildtype and Knockout groups and the percent of multipolar cells were quantified. **d, e** Representative Immunofluorescence staining images of bipolar (normal) and multipolar mitotic cells in wildtype and SETDB1 knockout PDC4 and 10 cell lines. Red: γ-tubulin, green: β-tubulin, blue: genomic DNA. Red arrowhead points to the abnormal mitotic cells. **f** Percent multipolar mitotic cells in wildtype and SETDB1 knockout cells in PDC4, 10 cell lines. (n=3 biological replicates) More than 25 mitotic cells in different fields were analyzed for abnormalities in each biological replicates within wildtype and knockout groups and the percent of multipolar cells were quantified. **g** Graphical scheme illustrating loss of SETDB1 results in loss of H3K9me3 and SETDB1 at peri-centromeric regions within different chromosomes. Knocking out SETDB1 can further promote abnormal cell division in the form of multipolar spindle formation during mitosis. **c, f**, Data shown as mean ± SD. P values were calculated using Unpaired t-test with Welsh’s correction.

Building on our findings in Ishikawa cell lines, we extended our experiments to two novel PDC cell lines (PDC4 and PDC10). Strikingly, imaging revealed the presence of multipolar cells in both cell lines (Fig. 5d, e, Supplementary Fig. 8c, d). Specifically, in PDC10, a recurrent primary EC cell line, there was about 38% abnormal cell division in the knockout cells verses about 18% of abnormal mitotic cells in wildtype (Fig. 5f). These findings highlighted the potential significance of SETDB1 governing proper cell division, shown in graphical schema (Fig. 5g), even though further studies of the mechanism are warranted.

### SETDB1 promotes immune evasion through blocking macrophage infiltration as well as promoting tumor proliferation

SETDB1^-/-^ tumors appear to grow at a significantly slower pace in mice compared to in cell culture. This led us to conclude that other external factors may contribute to tumor regression. Considering the importance of the immune system’s role in tumor growth, we conducted an analysis of the immune microenvironment within Ishikawa wildtype and SETDB1^-/-^ tumors. Since NSG mice are immune-compromised and lack key components of the adaptive immune system (ex. B-cells and T-cells) as well as NK cells, the innate immune system, primarily macrophages and granulocytes, constitutes the main cellular components present. Focusing on the CD11b+F4/80+ macrophage compartment in tumors, we observed a significantly higher presence of macrophages in SETDB1^-/-^ tumor than the wildtype by flow cytometry (Fig. 6a, b). Moreover, IHC staining of mouse tumor tissues with macrophage specific F4/80 antibody also showed greater infiltration of macrophages in SETDB1^-/-^ tumors (Fig. 6c). Further, mitotic figures assessment showed greater than 2-fold reduction in SETDB1^-/-^ tumors (Fig. 6d). Staining tumors with proliferation markers Ki67 and pHH3-ser10 confirmed reduced proliferation in SETDB1^-/-^ tumors (Fig. 6e-g). Overall, we demonstrated that SETDB1 promotes EC tumor progression through dual mechanisms: an extrinsic mechanism that represses immune cell infiltration into tumors and an intrinsic mechanism that promotes tumor growth.

**Fig. 6.**
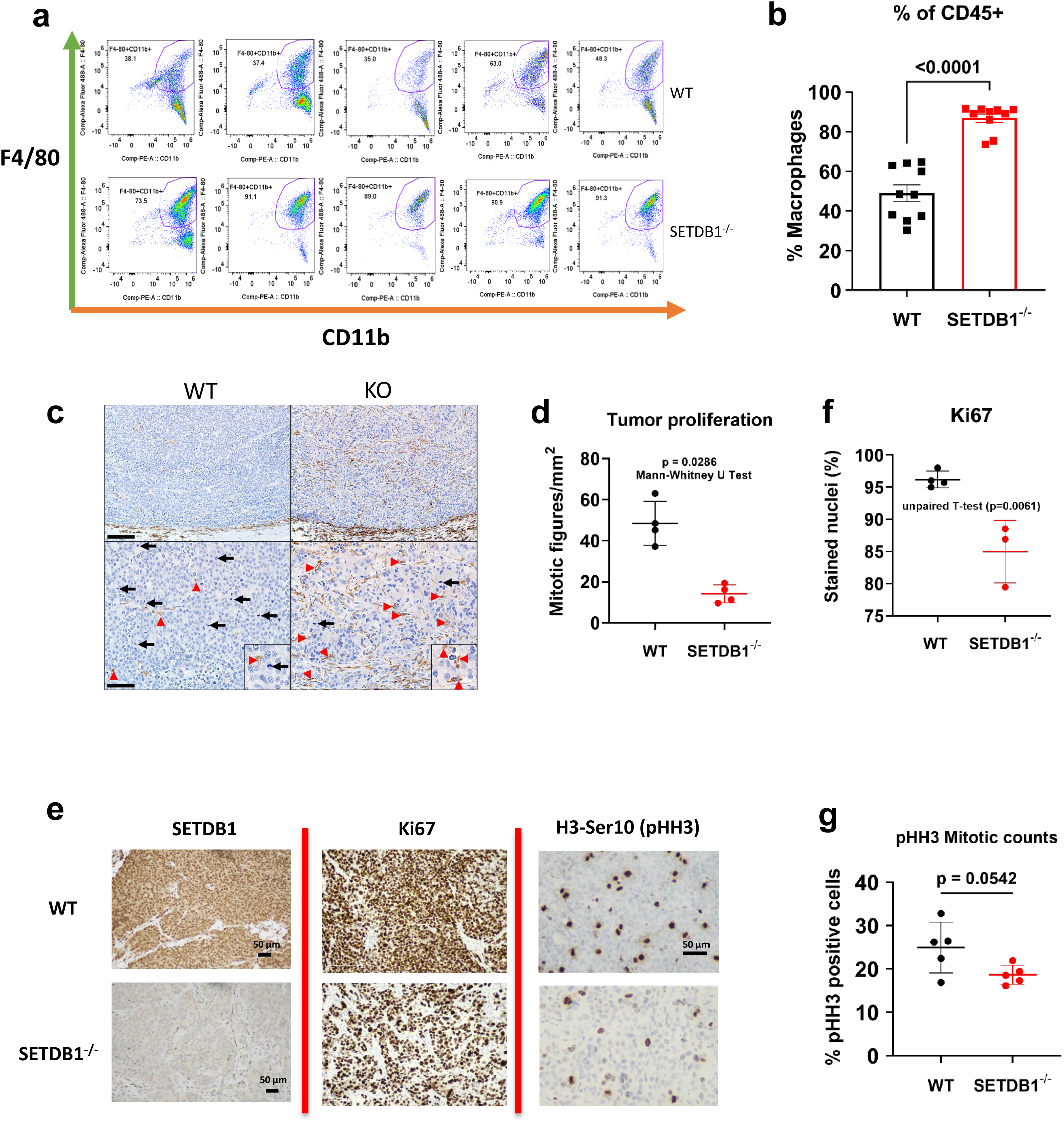
knockout SETDB1 promoted increased infiltration, recruitment of macrophages and decreased proliferation in tumor. **a** Representative gating strategy on CD11b+F4/80+ macrophages in wildtype and SETDB1^-/-^ Ishikawa tumors. The numbers on the plot represent the percent of macrophages from the total viable CD45+ cells. **b** Percent CD11b+F4/80+ macrophages quantification (out of the total CD45+ population) in wildtype and SETDB1^-/-^ Ishikawa tumors (n=10 per group). **c** Immunohistochemistry (IHC) staining for F4/80 on wildtype and SETDB1^-/-^ Ishikawa tumors. Red pointers represent brown stained F4/80+ macrophages and black pointers represent mitotic cells. Top and bottom scale bars represent 160 and 80μm respectively. **d** Quantification of mitotic figures per area (mm^2^) in wildtype and SETDB1^-/-^ Ishikawa tumors as an indicator of tumor proliferation (n=4 biological replicates). **e** Representative images of IHC stained wildtype and SETDB1^-/-^ Ishikawa tumors for SETDB1, Ki67, and phospho-H3S10 on histone (pHH3-ser10). **f** Percent measurement of Ki67 positive cells in wildtype and SETDB1^-/-^ Ishikawa tumors (n=4 per group). **g** Percent measurement of pHH3 positive cells in wildtype and SETDB1^-/-^ Ishikawa tumors (n=4 per group). Statistical tests: Student’s t-test (**b**) Mann-Whitney U test (**d**) Unpaired Student’s t-test (**f, g**). ns, not significant.

### Mechanisms of SETDB1 mediated immune escape: repression of repeat elements and interferon pathway

It has been reported that repeat elements and interferon pathways are crucial in activating innate immune systems. To investigate whether these elements and pathways are involved in macrophage recruitment, we analyzed the RNA expression of these genes in SETDB1^-/-^ tumors. We observed upregulation of all the endogenous retroviruses (ERVs), particularly, ERV class I and II (Supplementary Fig. 9a). Members of the interferon α and γ pathways such as TRIM21, PARP12, HLA-C, and CD40, showed increased expression in SETDB1^-/-^ cells and were among the most significantly altered genes (Supplementary Fig. 9b). Additionally, members of LINE (Long Interspersed Nuclear Elements) and SINE (Short Interspersed Nuclear Elements) exhibited significant elevation in SETDB1^-/-^ cells (Supplementary Fig. 9c, d). Collectively, these findings suggest that SETDB1 mediates immune escape through the repression of repeat elements and interferon pathways.

### SETDB1^-/-^ EC cells upregulate CCL5 expression in tumor and macrophage

The role of SETDB1 in regulation of immune related pathways in cancer has previously been well established. Given that we observed infiltration of macrophages in SETDB1^-/-^ tumors, we aimed to uncover a possible mechanism by which SETDB1 depletion leads to greater macrophage recruitment. We screened 17 mouse chemokines and found several such as *Cxcl10, Ccl3, Ccl4*, and *Ccl5*, showing significant upregulation in SETDB1^-/-^ tumors (Supplementary Fig. 10a, b). Notably, neutrophil-related chemokines, such as *Cxcl1* and *Cxcl2*, did not show significant alterations (Supplementary Fig. 10c), which aligns with our observation that SETDB1 knockout do not affect neutrophil infiltration in tumors. (Supplementary Fig. 10d, e).

It was reported that type I and II interferons show upregulation in SETDB1^-/-^ cancer cells such as in AML and in particular types of immune cells such as helper T cells^36, 37^. To identify human-specific cytokines that may impact macrophage recruitment by cancer cells, we performed qPCR. Among the cytokines tested, only *CCL5* showed the most upregulation in SETDB1^-/-^ tumors (Fig 7a). Considering the CCL5’s crucial function in promoting recruitment of immune cells (macrophages and T cells)^38^, we focused on CCL5 expression across different cells and tumors. We confirmed the upregulation of *CCL5*, as well as *CXCL9*, another important chemokine involved in immune cell recruitment, in our fixed timepoint mice SETDB1^-/-^ tumors (Fig. 7b).

**Fig. 7.**
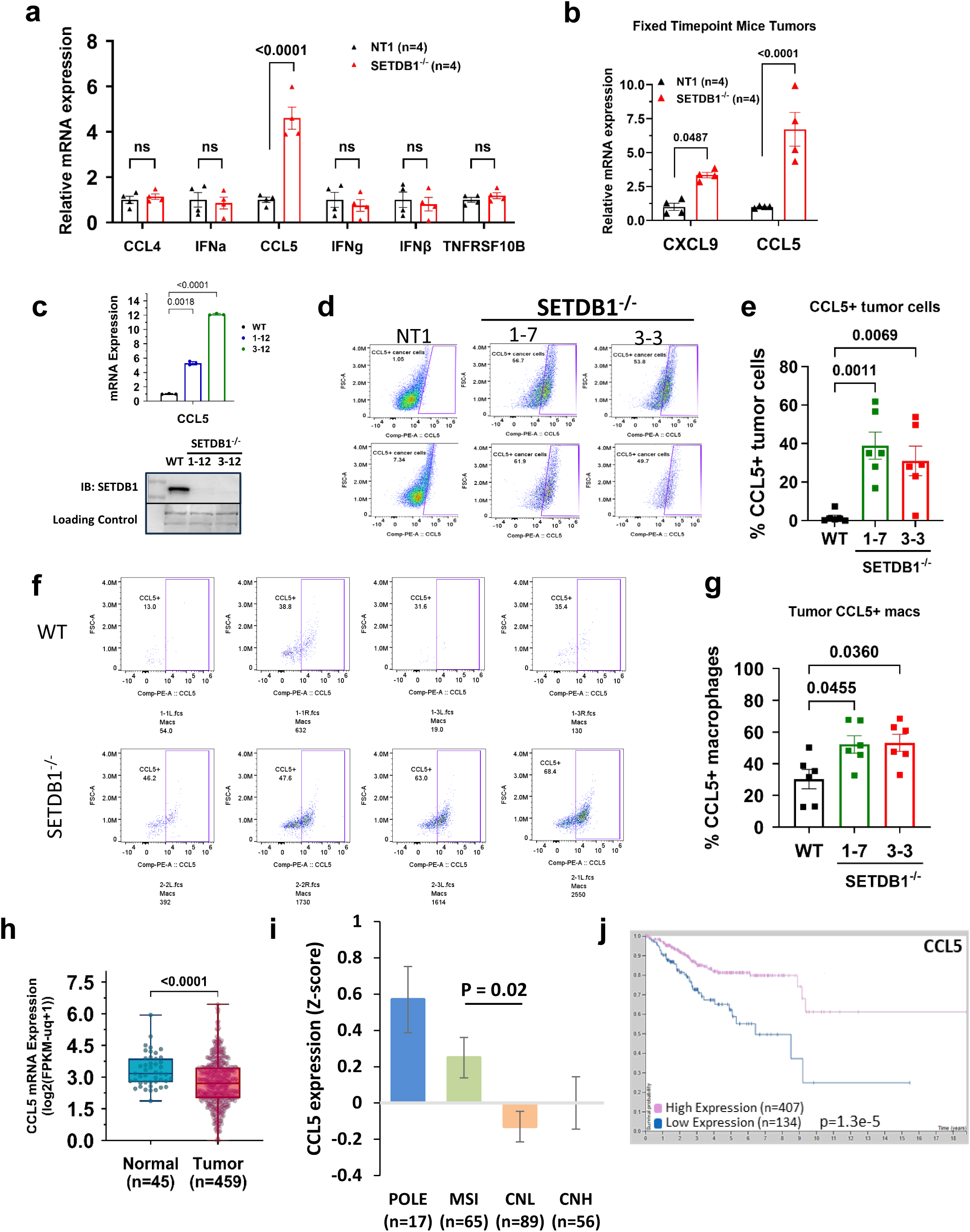
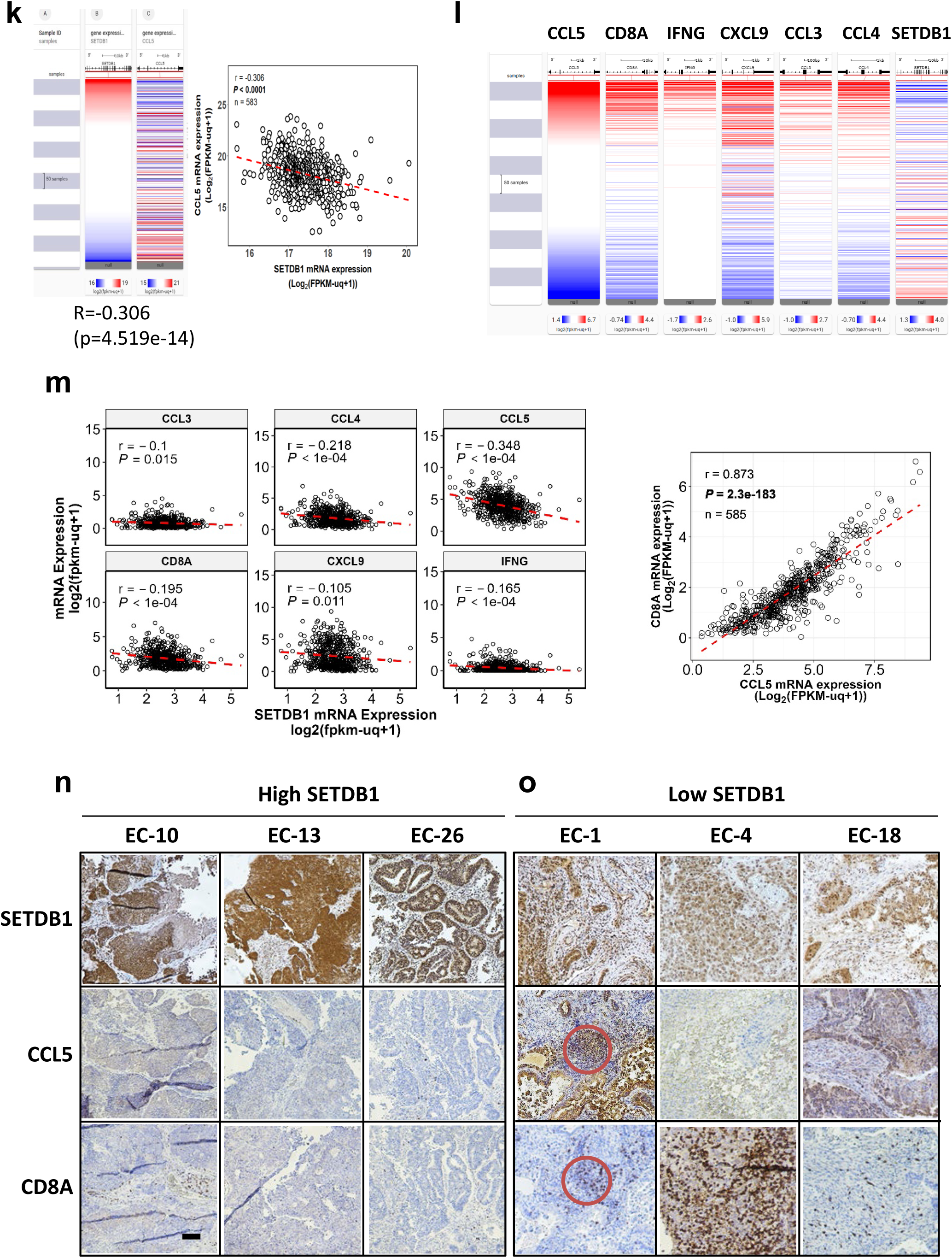
SETDB1^-/-^ EC cells upregulate CCL5 expression in tumor and macrophage. For **a-g**, all experiments performed in Ishikawa cells/tumors. **a** Quantification of human-specific cytokines and interferons mRNA expressions in SETDB1^-/-^ and wildtype tumors (n=4/group). **b** mRNA expression of CXCL9 and CCL5 in SETDB1^-/-^ and wildtype tumors in fixed timepoint tumor harvest (n=4/group). **c** Quantification of CCL5 mRNA expression in SETDB1^-/-^ and wildtype cell lines (n=3 technical replicates). **d** Representative gating strategy for CCL5+CD45-EpCAM+ wildtype, 1-7 and 3-3 SETDB1^-/-^ tumors. **e** Percent measurement of CCL5+ cancer cells in wildtype, 1-7 and 3-3 SETDB1^-/-^ tumors (n=6/group). **f** Representative gating strategy for percent CCL5+ macrophages in SETDB1^-/-^ and wildtype tumors. **g** Percent CCL5+ macrophages in SETDB1^-/-^ and wildtype tumors (n=6/group). **h** CCL5 mRNA expression across normal and tumor tissues in CPTAC dataset (Normal: n=45, and Tumor: n=459). **i** CCL5 mRNA expression across different EC subtypes. **j** Kaplan Meier curve illustrating patient prognosis for CCL5 high (n=407) and low (n=134) groups. **k** Heatmap and scatter plot displaying correlation for SETDB1 and CCL5 mRNA expressions utilizing EC-GDC TCGA dataset (n=583). **l, m** Heatmaps (**l**) and scatter plots (**m**) illustrating correlations for SETDB1 mRNA expression with T cell markers, and macrophage recruiting cytokines in EC-GDC TCGA dataset (n=583), respectively. Pearson correlation (r) and p values (*P*) are being displayed on the plot (n=583). **n**, **o** Images depicting IHC staining for SETDB1, CCL5, and CD8A on SETDB1 high and low expressing EC tumors respectively. Scale bar represents 100μm. For **c,** Data shown as mean ± SD. For **a, b, e, g, i**, Data shown as mean ± SEM. In **h**, boxplots display the full data range. The low and high ends of the box represent 25th and 75th percentiles, respectively. The middle horizontal line represents the median, with all data points displayed. Statistical tests: Two-way ANOVA (**a**, **b**), Welsh’s ANOVA (**c, g**), One-way ANOVA (**e, i**) with post hoc Sidak (**a**, **b**), Dunnett’s T3 (**c, g**), Dunnett’s (**e**), Bonferroni (**i**) tests. Welsh’s t-test (**h**) Pearson’s correlation coefficient (**m**). ns, not significant.

Like SETDB1^-/-^ tumors, CCL5 upregulation was confirmed in two SETDB1^-/-^ clones (Fig. 7c). This upregulation was also evident (up to 40% more) at the protein level when analyzing human EpCAM+CD45-cancer cells in SETDB1^-/-^ tumors through flow cytometry (Fig. 7d, e). Besides elevated CCL5 in SETDB1^-/-^ cancer cells, significant CCL5 upregulation was also evident in tumor associated CD11b+F4/80+ macrophages (Fig. 7f, g). These findings demonstrate that SETDB1 regulates CCL5 in EC.

To verify the clinical relevance, the CPTAC dataset was analyzed. CCL5 shows a significant decrease in RNA expression in tumor tissues (n=459) compared to normal tissues (n=45) (Fig. 7h). Intriguingly, among different molecular subtypes of EC, the CNH group of patients shows lower expression of CCL5 compared to POLE or MSI patients (Fig. 7i). This illustrates an inverse correlation with SETDB1 expression across subgroups (Fig. 1c). Consistently, data from the EC-TCGA dataset indicates that higher expression of CCL5 predicts better overall survival (Fig. 7j). CCL5 also has a negative correlation (R=-0.306, p=4.519e-14) with SETDB1 expression in EC-TCGA (Figure 7k).

Consistent with findings on mouse cytokines, SETDB1 demonstrated a reverse correlation with all upregulated cytokines such as CCL5 (r =-0.348, p < 0.0001), CCL3 (r =-0.1, p = 0.015), and CCL4 (r =-0.218, p < 0.0001) in the EC-TCGA dataset (Fig. 7l, m). Moreover, SETDB1 displayed a negative correlation with various T cell inflammatory markers such as CD8A (r =-0.195, p < 0.0001), IFNG (r =-0.165, p < 0.0001), and CXCL9 (r =-0.105, p = 0.011). To validate the reverse correlation between SETDB1 and CCL5/CD8A at the protein level, we selected two sets of EC patient samples. The high SETDB1 expression group exhibited low CCL5 and CD8A levels and vice versa (Fig. 7n, o). These data are consistent with the reverse correlation observed in the TCGA dataset, suggesting that SETDB1 facilitates the immune escape through downregulation of CCL5 and CD8A expression.

In conclusion, our data demonstrated that CCL5 is repressed by SETDB1 and potentially play a central role in macrophages recruitment in tumors.

### SETDB1 depleted cancer cells demonstrate greater sensitivity to M1-like macrophages *in vitro*

Next, we sought to find out whether infiltrating macrophages in SETDB1^-/-^ tumors directly kill the tumors cells. We isolated bone marrow from the female NSG mice and differentiated them into macrophages as previously described^39, 40^. Bone marrow derived macrophages were either polarized into M1-like macrophages by IFN-γ and LPS stimulation or kept in a non-activated neutral state (M0).

SETDB1^-/-^ or WT Ishikawa cells were cocultured with M1 or M0 macrophages for 24 hours. To determine whether M1-like macrophages induced greater killing of SETDB1^-/-^ cancer cells, we stained the cocultured macrophage/cancer cell mixture with CD45 antibody and a viability dye (Zombie), followed by flow cytometric analysis (steps shown in Fig. 8a). No significant changes in cell death were observed in WT cells in M0 and M1 conditions. However, in SETDB1^-/-^ cells, increased cell death was observed in the M1 group (Fig. 8b, c).

**Fig. 8.**
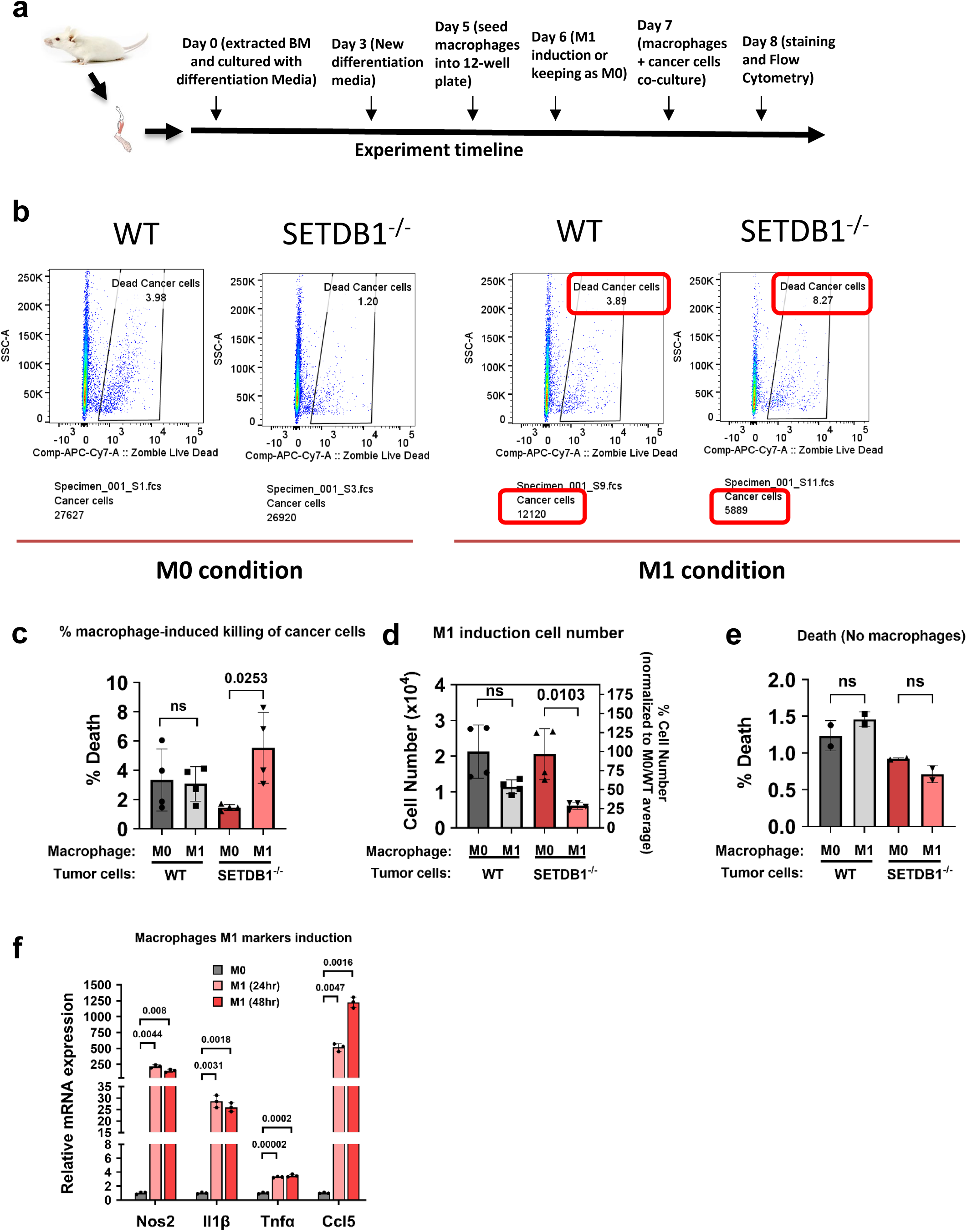
M1-like BM-derived macrophages induce more killing in SETDB1 deficient cancer cells in-vitro. **a** Experimental timeline for in-vitro BM-derived macrophage differentiation, M1 polarization, co-culture with cancer cells followed by flow cytometry staining and analysis. **b** Representative gating strategy measuring percent cancer cell deaths and changes in cell numbers induced by M1-like macrophages in co-culture treatment for WT and SETDB1^-/-^ Ishikawa cancer cells. **c** Percent death measurement of macrophage-induced cell death for wildtype and SETDB1^-/-^ Ishikawa cancer cells co-culture in M0 and M1 conditions (n=4 biological replicates). **d** Cell number quantification of wildtype and SETDB1^-/-^ cancer cells co-cultured with M0 or M1 induced macrophages (n=4 biological replicates). **e** Percent death measurement for wildtype and SETDB1^-/-^ cancer cells in M0 and M1 conditioned media (No macrophage present) (n=2 biological replicates). **f** qPCR mRNA quantification for M1 related markers Nos2, Il1β, Tnfα, and Ccl5 after macrophage M1 polarization with LPS and IFNγ for 24hr and 48hr (n=3 technical replicates). For **c-f,** Data shown as mean ± SD. Statistical tests: One-way ANOVA with post hoc Tukey’s multiple comparison test (**c,d,e**), Unpaired t-test with Welch correction (**f**). ns, not significant.

When measuring the total cell number, both WT and SETDB1^-/-^ cells showed a reduction when co-cultured with M1-like macrophages. However, the reduction was significantly higher for SETDB1^-/-^ cancer cells (Fig. 8d). These data suggest that SETDB1^-/-^ cells are more susceptible to macrophage-induced killing due to the reduced “don’t eat me” signaling. Additionally, M1-like macrophages conditioned media alone did not promote any cell killing in either WT or SETDB1^-/-^ cancer cells, indicating that cell death results from direct macrophage interaction (Fig. 8e). Indeed, when macrophages were polarized to M1, the M1 related genes such as Nos2, Il1β, Tnfα, and Ccl5 were all upregulated further proving that NSG macrophages are functional (Fig. 8f).

### Knocking out Setdb1 in mouse-derived EC cell line, MSH2-369, significantly reduces tumor growth and promotes macrophage infiltration in immunocompetent C57BL/6 mice

As NSG mice lack important adaptive immune system, to overcome this limitation, we extend our study in immune competent C57BL/6 mice. Mouse EC cell line MSH-369 cells (MSH2^-/-^ clone) were chosen and Setdb1 was successfully knocked out using LentiCRISPR-mediated knockout (Fig. 9a). Setdb1^-/-^ clones sg1-1, sg2-5, sg4-3 had significantly slower proliferation *in vitro* (Fig. 9b).

**Fig. 9.**
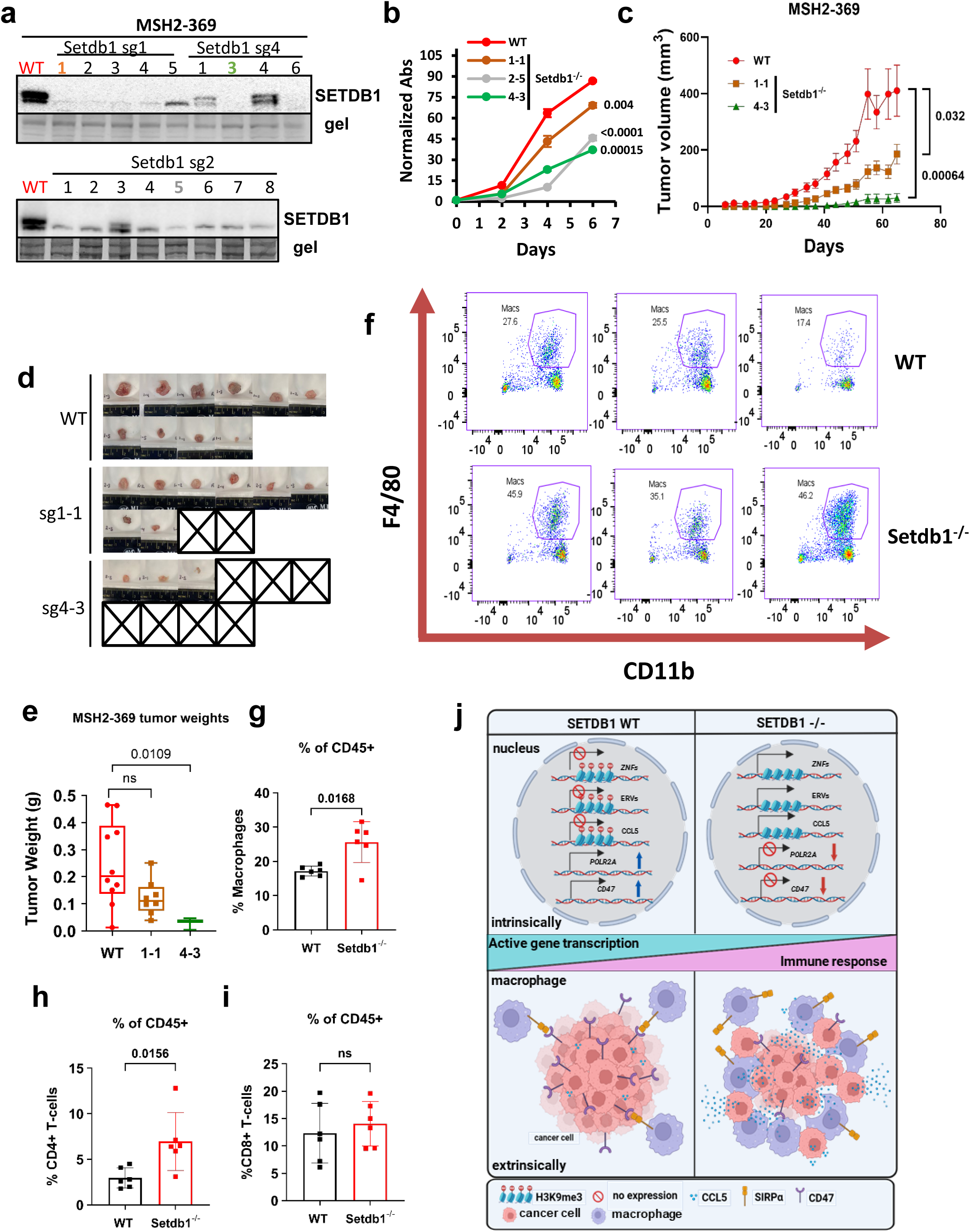
Knockout SETDB1 decreases tumor growth in immunocompetent mice (C57BL/6 tumor + immune cells). **a** Setdb1 Immunoblot for mice MSH2-369 Setdb1 knockout single clones from sg1 and sg4 Setdb1 lentiviruses. **b** Cell proliferation measurements for Setdb1 knockout clones and wildtype MSH2-369 cell lines (n=4 biological replicates). **c** Tumor growth trend for wildtype, 1-1, and 4-3 single clones of Setdb1 knockout subcutaneously injected in C57BL/6 mice (n=10/group). **d** Pictures of wildtype and Setdb1^-/-^ MSH2-369 tumors. **e** MSH2-369 tumor weights measurements for wildtype, 1-1, and 4-3 knockout clones (n=10/group for wildtype, n=8/group for 1-1, n=3/group for 4-3). **f** Gating strategy for measuring F4/80+CD11b+ macrophages in CD19-NK1.1-CD8-CD4-population subset for wildtype and Setdb1^-/-^ MSH2-369 tumors. **g** Percent of macrophages in total viable CD45+ cells in wildtype and Setdb1^-/-^ MSH2-369 tumors (paired t-test statistical analysis). For **b, g, h, i**, Data shown as mean ± SD. For **c**, Data shown as mean ± SEM. In **e**, boxplots have been spanned throughout the whole data. The low and high end of the box represent 25th and 75th percentiles respectively. Middle horizontal line represents the median. All the data points are visually displayed on the boxplot. **j** Graphical scheme shows dual functions of SETDB1: inside of nucleus, knock out SETDB1 promotes ZNFs, ERVs, CCL5, and inhibits POLR2A, CD47. in the cross talk between tumor cells and macrophages, knockout SETDB1 decreases CD47, advances CCL5 expression, which in turn attracts macrophages. Statistical tests: Unpaired Student’s t-tests (**b, c, g, h, i**) Kruskal-Wallis ANOVA test with post hoc Dunn’s multiple comparison test (**e**). ns, not significant.

Next, sg1-1 and sg4-3 knockout and wildtype cells were injected subcutaneously into C57BL/6 mice. Setdb1^-/-^ tumors exhibited a significantly decreased tumor growth rate. Strikingly, sg1-1 had 1/5 mice and sg4-3 had 3/5 mice with no tumor growth (Fig. 9c, d). Both knockouts had smaller tumor weights, however, only sg4-3 achieved statistical significance (Fig. 9e). Next, we explored how the immune system plays a role in tumor regression.

Using various immune cell markers (i.e. CD45, CD3, CD19, CD11b, F4/80, NK1.1, CD4+, CD8+, etc.), we sought to quantify different immune cells (i.e. T and B-lymphocytes, NK cells, and macrophages) and their proportions in the wildtype and Setdb1^-/-^ tumors. Intriguingly, we observed significantly more macrophages in Setdb1^-/-^ tumors (∼25.6% of the total CD45+ immune cells) compared to wildtype (∼17.1%) (Figure 9f, g). This finding was consistent with our previous observation on NSG mice growing human derived EC cells (shown in Fig. 6a, b). Further examination of T cells revealed that the CD4+ T cell population in Setdb1^-/-^ was 6.97% vs. 2.97% in wildtype (Fig. 9h). In contrast, CD8+ T-cells in SETDB1^-/-^ tumors showed a non-significant upregulation (Fig. 9i). Other immune cell subtypes were either not present (ex. B-cells) or did not change significantly between wildtype and Setdb1^-/-^ tumors.

## Discussion

Epigenetic dysregulation plays a crucial role in tumor progression. Many epigenetic modifiers, such as EZH2^41, 42^, LSD1^43, 44^, ASF1A^45^, and Sin3B^46^, have dual functions in promoting tumor progression and mediating immune evasion. SETDB1 has also been reported to possess dual functions, though not within a single system; however, our current studies reveal a novel dual function in EC. Our conclusion is supported by three lines of evidence: (1) In EC patient samples, high SETDB1 expression correlates with active gene transcription, while low SETDB1 correlates with an active immune response; (2) *In vivo* studies show that knockout of SETDB1 decreases mitotic figures and proliferation markers, while encouraging macrophage infiltration into tumors; (3) At the molecular level, SETDB1 promotes oncogene expression, inhibits tumor suppressors, and suppresses many immune response-related genes. Besides these novel findings, another important discovery is that SETDB1 governs proper cell division through binding to the SAR region in the centromere, implying that elevated SETDB1 may advance proper tumor cell division. Overall, our findings reveal a novel mechanism by which oncogene SETDB1 drives tumor progression and facilitates immune evasion.

Though SETDB1 has been studied in more than 10 tumor types^4, 5^, our study is the first to focus on the function of SETDB1 in EC. Two studies have reported SETDB1 overexpression in EC^16, 17^; however, neither study explored the molecular mechanisms regulated by SETDB1. Among the four EC subtypes, the CNH/p53abn subtype is the most aggressive and poorly immunogenic. We found that SETDB1 expression is highest in this group. SETDB1’s dual functions promoting tumor growth intrinsically and mediating immune evasion extrinsically, are potential mechanisms contributing to the aggressive and poorly immunogenic nature of this group. This implies that SETDB1 is a driver gene in this subtype and may serve as a potential target.

Overall, we demonstrated SETDB1’s dual role in promoting cell proliferation and suppressing the anti-tumor immune response. We found several tumor suppressor genes were repressed by SETDB1 such as *RERG, PGR* and *ZNF582* which may play a role in impeding tumor growth in EC. PGR is a well-documented tumor suppressor in EC^20^. RERG has been reported to suppress tumor growth in nasopharyngeal carcinoma and breast cancer through the suppression of ERK/NFκB signaling and inhibiting anchorage independent growth, respectively^21, 23^. Furthermore, ZNF582 impairs the progression of clear cell renal cell carcinoma by inhibiting ERK2 phosphorylation and suppressing the MAP kinase signaling pathway^25^. Additionally, hypermethylation of ZNF582 promotes metastasis of nasopharyngeal carcinoma by regulating the transcription of Nectin3 and NRXN3^47^. A variety of known oncogenes have shown direct expression correlation with SETDB1, such as *FLNA*, *POLR2A,* and *MUC5AC*. FLNA promotes tumor growth by activating the PI3K/AKT pathway in melanoma through interaction with TRIP13^48^ and by positively regulating ERK and AKT activation in KRAS-induced lung tumors^49^. POLR2A synthesizes mRNA, which is critical for controlling cell proliferation, such as in gastric cancer and triple-negative breast cancer^27, 50^. MUC5AC’s role is promoting cancer cell stemness and chemoresistance in pancreatic cancer ^51, 52^, and is essential in KRAS-dependent lung cancer tumorigenesis^53^. Altogether, we confirmed multiple SETDB1 target genes, elucidating SETDB1’s intrinsic function in tumorigenesis and potentially serving as an activity signature.

Gene signatures have been well explored by both bioinformatics and bench scientists across different cancers to predict survival and drug sensitivity^54, 55, 56^. Specifically, in EC, multiple signatures are suggested to facilitate the prognosis, as well as distinguish the stage and recurrence^57, 58, 59^. For example, utilizing the TCGA dataset on UCEC, two studies predicted overall EC survival using a six or four gene signature^60, 61^. However, these studies lacked experimental validation. In contrast, we found that SETDB1 activity signature “SETDB1 + MSH6 – PGR” distinguishes tumor grade, while another SETDBI activity signature “SETDB1 + CD47 – PGR” predicts patient survival. Thus, our findings conclude that SETDB1 not only impacts tumorigenesis in EC, but also predicts both tumor grades and patient prognostic outcomes.

For the first time, we report a novel and unexpected role for SETDB1: it binds and promotes H3K9me3 at SAR regions within different chromosomes in human cancer cells, as revealed by our ChIP-seq analysis. While H3K9me3 has been reported to bind centromeres, governing chromatin accessibility and chromosomal stability ^62^, no specific methyltransferase has been identified, nor detailed mechanism or outcome revealed. Over the past years, SARs have been shown to play a role in many central cellular processes ranging from DNA replication, DNA repair, to transcription and RNA processing^63^. Our studies revealed that the loss of SETDB1 leads to abnormal cell division, which is consistent with studies in mouse germlines, but novel in human cancer cells^35^. Although further studies are needed to reveal the mechanism behind this finding, this highlights the importance of SETDB1 as a cell division regulator. Chromosomal missegregation can lead to chromosomal instability (CIN) and possibly cell death^64^, which may explain the observed lower tumor proliferation upon SETDB1 loss.

We are the first group to report that beyond the mechanistic role in tumorigenesis, SETDB1 establishes immune evasion in EC. Several previous publications have identified SETDB1 as an important mediator of immune response. One outstanding study marked SETDB1 as the top candidate among 936 epigenetic modulators to induce immunotherapy resistance^6^. We discovered five mechanisms involved in SETDB1-mediated immune evasion through: (1) promoting the “don’t-eat-me” signal, CD47; (2) blocking macrophage infiltration, (3) inhibiting ERVs and IFN pathways; (4) decreasing macrophage and T-cell chemokines, CCL5 and CXCL9; (5) preventing CD4+ T-cell infiltration.

First, although CD47 expression has been reported to be regulated by HIF1a, MYC, and cytokine signaling^28, 65,67^, we made a novel discovery that SETDB1 is able to promote CD47 expression. CD47 is known for its role as a “don’t-eat-me” signal which often is overexpressed in multiple cancers^28, 66^, including EC^67^. Through binding with its interacting partner, Signal Regulatory Protein alpha (SIRPa) on the surface of macrophages, it can block any phagocytic activity against the tumor cells^68^. Reduction in CD47 expression on SETDB1^-/-^ cells means that they have higher vulnerabilities to macrophage mediated phagocytosis and immune exclusion.

Second, we are the first group to report that knocking out SETDB1 encourages macrophage infiltration. Traditionally, macrophages in NSG mice are considered defective, however, our studies demonstrated their functionality and ability to target tumor cells^69, 70, 71, 72, 73, 74^. When cocultured, the M1 polarized bone marrow-derived macrophages from NSG mice kill 70% of SETDB1^-/-^ cells verses only 40% of WT cells (Fig. 8d). The macrophages infiltrated the SETDB1^-/-^ cells more than the WT, limiting the killing effect to the WT, but enhancing the killing effect on SETDB1^-/-^ tumors (Fig. 6a, c). Indeed, M1 markers such as CCL2 and NOS2 mRNA expression increased upon M1 polarization (Fig. 8f). Additionally, we confirmed macrophage infiltration in immunocompetent C57BL/6 mice, where macrophages were recruited into the SETDB1^-/-^ tumor.

Third, it was reported that the downregulation of type I interferon signaling pathway is central to SETDB1-mediated immune escape in ovarian cancer, AML, and melanoma^36, 75, 76, 77^. Specifically, the mechanism involves SETDB1 depletion, promoting the induction of interferon response genes (ISGs) through the activation of retrotransposons and ERVs. Our studies are consistent with the report; we verified that the loss of SETDB1 resulted in enhanced production of ERVs and both type I and type II IFN responses.

Fourth, we showed that SETDB1 hindered macrophage infiltration in tumors, possibly by blocking the expression and secretion of CCL5. Consistent with other reports^78^, we discovered that SETDB1 represses chemokine CCL5 expression. CCL5 cytokine release by tumor has been demonstrated to be crucial for CD8+ T-cell infiltration. Dangaj et al reported a direct correlation between tumor cells that expressed CCL5 and CD8+ T-cell presence^38^, where CCL5 expression stimulates T-cells to produce IFN-y, promoting monocytes/DCs (dendritic cells). This produces CXCL9 which encourages CXCR3+ CD8+ T-cells to infiltrate the tumor. Thus, CXCL9 and CCL5 are important components in developing T-cell-mediated anti-tumor responses. Consistently, we validated the increase in expression of both CCL5 and CXCL9 in our SETDB1^-/-^ tumors. In the TCGA dataset, the mRNA expression of CCL5 and CD8A showed a strong positive correlation (r=0.873, p=2.3e-183) (Fig. 7o). Next, we were motivated to check CD8A infiltration in SETDB1^-/-^ tumors.

Last, previous publications have presented SETDB1 as a key regulator of CD8+ cytotoxic T cell anti-tumor function^77, 78^. Intriguingly, we observed a slight increase in CD8+ T cells in SETDB1^-/-^ tumors in C57BL/6 mice, possibly due to three possible reasons: (1) MSH2^-/-^ tumor already has enhanced immunogeneicity due to MSH2 being an important MMR player, (2) failure to collect the CD8+ T cell clusters during tumor pieces staining, and (3) limited sample size. Therefore, we did not observe a further boost in CD8+ T cells in MSH2 Setdb1^-/-^ tumors. In the future, more mice need to be injected with tumors to expand the sample size. Additionally, less immunogenic mice EC models, such as MECPK tumors (Mouse EC Pten-/-Kras G12D knock-in), may represent an alternative system to evluate the effect of SETDB1-mediated immune response. Aside from these limitations, TCGA data showed a reverse correlation between SETDB1 and CD8A, and our IHC staining confirmed increased CCL5 and CD8A in low SETDB1 expressing tumors. Furthermore, we observed an inverse correlation between SETDB1 and the mRNA level of CCL5, CXCL9, CD8A in TCGA. These findings suggest that SETDB1 could serve as a target for immunotherapy to direct CD8+ T-cells against tumor in the clinical setting.

We acknowledge there are limitations in our studies. First, we aim to verify whether depleting macrophages will rescue SETDB1-/-tumor growth. Clodronate liposome treatment was a well-accepted strategy to deplete macrophages, however, due to toxicity of the clodronate in NSG mice, we did not achieve positive results. In the future, this toxicity may be solved by different resources of chlodronate liposome or different mice models. Alternatively, we can use macrophage-null mouse model Csf1r-/-to verify the crucial function of macrophage in limiting tumor growth. Second, more EC tumor models will be important to generalize our findings regarding SETDB1’s critical function in EC.

We recognize that further studies are needed to fully understand SETDB1 functions in cancer. On the basic science level, investigating how SETDB1 supports the interaction of the kinetochore with the mitotic spindle and facilitates cell division will further elucidate its critical functions. Given that SETDB1-mediated cell division has been reported in mouse germlines, we hypothesize that SETDB1’s role in proper cell division is conserved across species and can be generalized to other cancer types.

However, further studies are warranted. To confirm that CD47 is the critical factor for SETDB1-mediated immune evasion, CD47 neutralization antibodies can be utilized to prevent macrophage phagocytosis. To validate whether CCL5 is a critical factor in recruiting macrophage and T cells in SETDB1-/-tumors, potential strategies using neutralization antibodies for CCL5 may prevent macrophage and T-cell infiltration. On the translational level, supported by pan-cancer studies, SETDB1 expression is inversely correlated with immune response pathways and positively correlated with cell proliferation pathways. We hypothesize that dual oncogenic functions of SETDB1 can be applied to other cancer types, in which SETDB1 is overexpressed and negatively impact patient outcomes.

In this study, we elucidated the multifaceted role of SETDB1 in EC growth and its impact on the tumor microenvironment. Our findings highlight that the SETDB1 activity signature is crucial for EC diagnosis and prognosis, as its overexpression is associated with worse survival outcomes. Targeting SETDB1 in EC may be especially useful in the most aggressive and poorly immunogenic CNH/p53abn group, which exhibits the highest SETDB1 expression among the subtypes. Inhibiting SETDB1 enhances immunotherapy sensitivity by promoting tumor antigen expression and increasing T-cell infiltration.

Additionally, SETDB1 inhibition could boost antitumor immunity. Our novel findings suggest that targeting SETDB1 could be a promising therapeutic strategy not only for EC but also for other cancers with elevated SETDB1 expression, thus providing a strong rationale for the development of SETDB1 inhibitors as potential adjuncts to existing cancer therapies.

## Methods

### Animals

All animal related experiments were approved by Broad Institution of Animal Care and Use Committee (IACUC). Animals were kept at University of Iowa Animal facility. NOD.Cg-Prkdc^scid^ Il2rg^tm1Wjl^/SzJ (NSG) (stock number: 005557) and C57BL/6J (stock number: 000664) mice were purchased from Jackson Laboratory. Mice were maintained and propagated under IACUC and institutional guidelines under pathogen-free conditions.

### Cell lines

Ishikawa, ECC1, and Hec50 human EC cell lines were obtained from ATCC. MSH2-369 is a murine EC cell line^79^ kindly provided by Dr. Melinda Yates (University of North Carolina, Chapel Hill, NC, 27599, USA). For PDC cell lines (ex. PDC4, 8 and 10), patient consent was acquired with IRB approval. All cells are free of mycoplasma, as detected every six months using MycoStrip (Mycoplasma Detection Kit, InvivoGene, cat#: rep-mys-50). Tumors were collected from operating room, then implanted in NSG mice. At endpoint, tumors were removed, digested, and cultured in cell culture dish. They were cultured in DMEM medium (Ishikawa, Hec50), RPMI1640 medium (ECC1), or DMEM/F12 medium (MSH2-369, PDC4, PDC8, and PDC10) supplemented 5% FBS and 1% penicillin/streptomycin at 37℃ in a humidified atmosphere with 5% CO_2_.

### In-vitro cell proliferation Assay

For Ishikawa, ECC1, and Hec50, cell proliferation rate was measured by plating cells in 6-well plate at 100,000 cells in triplicates. At 3, 5, and 7 days timepoints, cells were trypsinized. Automatic cell counter was used to count the cells in each well. The number of cells were then averaged for the three cell count readings at each timepoint, standard deviation was calculated, and the growth curve was produced using Microsoft Excel. For PDCs and MSH2-369 cell lines, the proliferation was measured using resazurin assay. Briefly, resazurin was added to cells cultured in 96-well plate at day 0 (when cells were just seeded) and other timepoints such as day 4 (PDC8 and 10) and day 10 (PDC4) after culture.

Fluorescence was measured (excitation: 530nm, emission: 590nm), absorbance values were normalized against day 0 average reading, standard deviation was calculated. Average values for the desired timepoints for SETDB1 ko cells were then normalized against wildtype as % growth and illustrated as bar plot. For MSH2-369, absorbance readings for each timepoint were normalized against day 0 average, standard deviation was calculated and plotted as growth curve.

### Tumor models

All tumors were established by injecting the cancer cells subcutaneously into left and right upper abdomen of the female mice. Ishikawa, ECC1 and Hec50 tumors were grown in NSG, and MSH2-369 tumor was developed in C57BL/6 mice. In the 1^st^ injections (for Ishikawa and ECC1), 1×10^6^ and 2×10^6^ cells were injected for Ishikawa and ECC1 cell lines respectively. In the survival experiments, 1×10^6^ cells were injected for Ishikawa and ECC1, and 0.5×10^6^ cells were injected for Hec50 cell line. As for the fixed time point experiment, 5×10^6^ Ishikawa cells were injected. For MSH2-369 cell line, 1×10^6^ cells were injected. MSH2-369 cells were established from endometrial tumors derived from PR-Cre *Msh2*^flox/flox^ (abbreviated Msh2^-/-^) C57BL/6 mouse^79^.

### Quantitative Real-Time PCR

After both tumor and cell lines mRNA extracted using Monarch Total RNA Miniprep Kit (NEB #T2110) and purified, mRNA concentration was measured using Nanodrop. Then, mRNA was reverse transcribed to cDNA using BioLabs Protoscript II First Strand cDNA Synthesis Kit (Cat# E6560L). cDNA was used for quantitative PCR with the PowerUp SYBR Green PCR Master Mix (Applied Biosystems, A25742). Signals from PCR was detected using QuantStudio3 (Applied Biosystems). The sequences of the PCR primers are listed in Supplementary table 1. The Ct values were acquired for each primer and samples. GAPDH was used as internal control gene. For each sample, first, the GAPDH Ct value was subtracted from the Ct value of the gene of interest (GOI) (ΔCt). The Fold Change (FC) was calculated using the formula 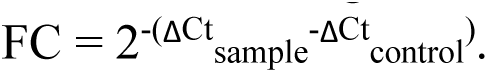

### Western blotting

Cell lines were washed twice in cold PBS. Cell Lysates were then prepared using 50mM Tris pH7.2, 150mM NaCl, 1% NP-40 solution supplemented with Proteinase Inhibitor and phosphatase Inhibitor (ThermoFisher, Cat# 78443). Protein concentration of the lysates was then measured using Pierce protein assay BCA kit (Cat# 23225). Tumors were lysed using RIPA buffer (10mM Tris pH 8, 1mM EDTA, 0.5mM EGTA, 1% Triton X-100, 0.1% Sodium Deoxycholate, 0.1% SDS, 140mM NaCl).

Protein extracts were run and separated on 7.5-12% SDS-PAGE gels, transferred to Nitrocellulose membrane, blocked with 5% Milk in Tris-buffered saline (25mM Tris pH7.2, 135mM NaCl) supplemented with 0.1% Tween-20 (TBST). Membranes were then incubated at 4℃ overnight with the primary antibodies SETDB1 (Cell Signaling #93212S, #C1C12, ProteinTech, Cat# 11231-1-AP) diluted in 5% BSA in TBST. Next day, membranes were washed twice in TBST and incubated 2hr in HRP anti-rabbit secondary antibody diluted 1:10,000 in 5% Milk-TBST. Next, membranes were washed three times with TBST. Membranes were then treated with ECL reagents to produce luminescence signal, and the signal was read by Chemidoc MP imaging system (BioRad).

### Bone Marrow derived macrophages and killing assay

Bone marrow was extracted from NSG mice femur and tibia bone. The bones were cut at both ends and using a 5ml syringe containing PBS + 25ug/ml Gentamycin and 26G1/2 needle, the bone marrow was flushed into 15ml tube. Each mouse isolated bone marrow was then incubated into two petri dishes and are differentiated into macrophages using differentiation media (RPMI 1640 + 10%FBS + Pen/Strep+ 25ug/ml Gentamycin + 20ng/ml M-CSF) for a total of 6 days. Media was changed and renewed 3 days after the start of differentiation. On day 5, macrophages were removed from the plate by scraping and plated in 12-well plate for preparation of co-culture with cancer cells. Either 100,000 (for 1:1 ratio) or 300,000 (for 1:3 ratio) macrophages were plated. On day 6, macrophages are either polarized into M1 using M1 media (Differentiation media + 100ng/ml LPS + 20ng/ml IFNg) or maintained at the M0 state by differentiation Media alone. On day 7, cancer cells (either control or SETDB1^-/-^ cells) co-cultured at 100,000 cells in each well. Next day, co-cultured cells were treated with trypsin followed by scraping to harvest macrophage cancer cells mix. The mix were then stained with Zombie NIR viability dye (Biolegend #423105), anti-CD45 Alexa Fluor 700 (Biolegend #103127), anti-F4/80-Alexa Fluor 488 (Biolegend #123120) and anti-CD11b PE (Biolegend #101208) to quantify %dead cells and total viable cancer cells in the co-culture treatments using Cytek Aurora. FlowJo software was used to do the analysis.

### RNA-Sequencing and analysis

For whole RNA sequencing, Ishikawa wildtype and SETDB1 knockout single clones (1-7, 2-6, and 3-3) were cultured in triplicates in 100mm cell culture dishes. mRNA was extracted as mentioned earlier in method. Sample mRNAs was sent for quality verification with the Bioanalyzer (Agilent model 2100) and subsequent library preparation to the University of Iowa Genomics Division of the Iowa Institute of Human Genetics (IIHG). RNA sequencing (paired end reads,150-bp) was performed in the IIHG Core Facility using the Illumina NovaSeq. Quality control of the resulting fastq files was assessed using the FastQC program (version 0.11.5). Reads were mapped against the human reference genome (version hg38) using the STAR aligner (version 2.5.3a) and a file containing the counts mapping to each gene loaded into R (version 3.5.2). The raw counts were normalized and transformed using the rlog function. For statistical analysis of the data, the DESeq2 package (version 1.22.2) in R was used. In brief, a model incorporating all the experimental factors was created and Wald tests were used to compute statistical metrics. A gene was considered to have a statistically significant change in expression if the False Discovery Rate was less than 10%. Results from the statistical analysis were visualized using heatmaps (ComplexHeatmap, version 3.20) and volcano plots (ggplot2, version 3.4.4).

### Chromatin Immunoprecipitation (ChIP)

Cell lines were cultured in TC100 plates. Cell signaling SimpleChIP Enzymatic Chromatin IP Kit (#9003) and their ChIP protocol was used to conduct the ChIP. Briefly, cells were fixed with 0.1% paraformaldehyde for 10mins on shaker at room temperature. Glycine was added to quench the paraformaldehyde and was shaken for 5mins. Cells were washed afterward with 10ml of cold PBS twice. 1ml of PBS/protease inhibitor was added on top of the cells and cells were scraped off the plate and collected. Buffers A and B were added to cell pellets in order after centrifuging the cells to isolate the nucleus and chromatin. Micrococcal nuclease was used to cut the chromatin into small pieces for 20mins at 37℃ followed by inactivation with EDTA. Digested chromatin was then further sonicated for 3×20s intervals with 30s rest in between using Model 505 Sonic Dismembrator. After centrifuging for 10mins, a small sample of supernatant was taken to verify chromatin digestion and concentration through treatment with proteinase k and RNase A followed by purification, agarose gel run, and nanodrop measurement. 10ug of chromatin was used to conduct ChIP using anti-H3K9me3 antibody (cell signaling #13969, 1:100), anti-SETDB1 antibody (proteintech, 11231-1-AP, 1:167), anti-rabbit IgG (cell signaling #2729, 1:1000) overnight at 4℃. 4% of input was put aside for reference. The next day, protein G magnet beads were added to antibody-chromatin mix and incubated for 2hours with constant rotation at 4℃. Beads were then washed with salt solutions. Elution of the chromatin from antibody/beads were conducted in elution buffer at 65℃ for 30mins with constant rotation. Reverse cross-linking was then performed with proteinase k at 65℃ for 2hours. Final DNA fragments were eluted and purified using DNA purification spin column.

### ChIP-sequencing and analysis

Sequencing (50-bp paired end reads) was performed using the Illumina NovaSeq 6000 sequencer. We then used a custom computational pipeline developed with Nextflow (https://www.nextflow.io) to analyze the ChIP-Seq data. Quality of the reads was assessed using FastQC (version v0.11.5) and ngmerge (version 0.3) was used for read trimming. Trimmed reads were aligned to the human genome (build hg38) using the bwa mem aligner (version 0.7.15-r1140) and peak calling was performed with Genrich (version 0.5) using the following filtering options: removal of PCR duplicates and retention of unpaired alignments. The default p-value of 0.01 was used for statistical significance. Processed ChIP-Seq data containing peak location information (narrowPeak format) were imported into R (version 4.0.0) and analyzed using the ChIPseeker package (version 1.18.0). Visualization of peaks was performed using the Integrated Genomics Viewer (IGV version 2.11.2). To identify motifs in selected peaks, we first extracted peak sequences using the Biostrings package (version 2.50.2) in R followed by enrichment analysis with MEME (online version 5.5.7).

### Mice dissection, tumor dissociation and staining

For tissue collections, mice were sacrificed. Tissues were collected on ice in PBS. Tumor tissues were dissociated into single cells by first mincing the tumors into small pieces. Then, dissociation buffer (RPMI 1640 + 5% FBS + 0.5mg/ml collagenase A+ 0.02mg/ml DNase I) was used to dissociate tumors further for 45 min at 37℃. Dissociated tumors were then passed through 70μm cell strainer. Dissociated tumor was then washed in PBS. For staining, first dissociated tissue was stained with Zombie viability dye (1:1000) in PBS for 10-15mins. After PBS wash, surface marker antibodies Alexa Fluor 700 anti-Mouse CD45 (Biolegend #103127) 1:400, anti-Mouse F4/80-Alexa Fluor 488 (Biolegend #123120) 1:200, Brilliant Violet 650 anti-Mouse CD11b (Biolegend #101259) 1:500, Brilliant Violet 785 anti-Human EpCAM (Biolegend #324237) 1:50, BD-OptiBuild^TM^ BUV395 Rat anti-Mouse CD3 (BD Biosciences #740268) 1:100, Brilliant Violet 711 anti-Mouse CD8a (Biolegend #100747) 1:100, and BD Horizon BUV496 Rat anti-Mouse CD4 (BD Biosciences #612952) 1:100 were diluted in PBS and used to stain the tissues on ice for 20mins. Next, tissues were washed with PBS. For intracellular staining (ICS), the fixation and permeability kit from BD Biosciences was utilized to fix and permeabilize cells for ICS following their protocol. PE/Cy7 anti-CCL5 (Biolegend #149105) 1:200 was then diluted in permeabilization buffer and used to stain the cells for 20mins on ice. Next, cells were washed with permeabilization buffer and flow cytometry was conducted using Cytek Aurora.

### Immunohistochemistry (IHC) staining

Tissues were fixed, embedded, sectioned and stained in the Histology Research Laboratory at the University of Iowa. Briefly, tissues were fixed in 10% Neutral-buffered formalin (NBF) for 24-48hr, embedded in paraffin, and were cut at 5μm thickness. Tissues were mounted on microscope slides followed by immunohistochemistry staining on an Epredia Autostainer 360 with the UltraVision Quanto Detection System HRP DAB kit. Briefly, slides were dewaxed and antigen retrieval was performed using Epredia’s Dewax and HIER buffer L (#TA999DHBL) for 20 minutes at 95C. Slides were then treated with kit reagents (10 minutes for UVQuanto Endogenous Block, 5 minutes for UVQuanto Protein Block, primary antibody, 10 minutes with UVQuanto Amplifier, 10 minutes with UVQuanto Micro Polymer HRP, and 5 minutes with UVQuanto DAB). Primary antibody was used for 30 minutes for each of the following antibodies and concentrations: anti-F4/80 (#MCA497GA, Bio-Rad, 1:250), anti-pHH3-ser10 (#369-A14, Cell Marque, 1:200), ki67 (cat# ab16667, Abcam, 1:500), SETDB1 (cat#11231-1-AP, Protein tech, 1:200), CCL5 (cat# BS-1324R, Bioss, 1:200), CD8A (cat# 372902, Biolegend, 1:100). Counterstaining was performed with on a Leica Autostainer XL using Lecia Harris Hematoxylin (Cat. No. 3801560) for 3 minutes before clearing in alcohol and xylene. Slides were mounted and coverslipped with Permaslip.

### Immunofluorescence staining

Cells attached to coverslips were initially washed with PBS twice. Next, cold-20℃ Methanol was added on top of the cells for fixation. The cells were then incubated in-20℃ freezer for 15mins. PBS was then used to wash off methanol for three times. 5% BSA-PBS solution was used to promote blocking for 30 mins. Dilution for the primary antibodies β-Tubulin (Cell Signaling #15115) and γ-Tubulin (Sigma #T6557) were prepared at 1:100 and 1:1000 respectively. Primary antibodies were incubated with the fixed cells overnight at 4℃ in a humidified environment. On Day 2, coverslips were rinsed with PBS three times. Secondary antibodies Goat anti-Rabbit Alexa Fluor^TM^ 488 Plus (ThermoFischer #A32731) and Goat anti-Mouse Alexa Fluor^TM^ Plus 555 (ThermoFischer #A32727) were diluted 1:200 in 5%-PBS. The antibody solution was incubated on coverslips for 2 hours at room temperature. Coverslips were rinsed with PBS 5 times. Finally, the ProLong(R) Gold Antifade with DAPI (Cell Signaling #8961S) was utilized to mount the coverslips on the slide. Images were acquired by Nikon confocal *Ti2* and Eclipse Ci fluorescent microscopes.

### Plasmids and vectors

For knocking out SETDB1 in cancer cells, lentiCRISPR-Cas9 vector were used to generate the virus^80^. For overexpression, Lentiviral pLKO.1-SETDB1 vector was provided by Dr. Phillip Koeffler, Jonsson Comprehensive Cancer Center, UCLA. Mouse specific Setdb1 sgRNA sequences were obtained from Griffin et al., Nature, 2021 and Burbage et al., Sci Adv., 2023 papers. The SETDB1 sgRNA sequences are provided in Supplementary Table 2.

### Lentiviral packaging and transduction

For packaging lentiviruses, LentiCRISPRv1 vector containing the sgRNAs and Cas9 were mixed (2µg) with 1.5µg of psPAX2 (packaging) and 1µg of pVSV-G (envelope) in Opti-MEM Reduced I Serum media and then added to the 293T cells in 6-well plate. Cells were then passaged into poly-D-Lysine pre-coated 100mm cell culture dish in order for cells to produce the lentiviruses. 72hr after transfection, lentiviruses were collected in the media on top of the cells. Media containing the lentivirus was filtered through 0.45µm filter to exclude the 293T cells.

For transduction, target cells were grown in 6-well plates. Then, 2ml of the lentivirus added to 1ml of media on top of the target cells. 24hr after the transduction, target cells were passaged into 100mm plates and the day after were selected on 2µg/ml puromycin to select for transduced cells.

## Data availability

Publicly available datasets presented in this paper are from UCEC Xena Browser (GDC-TCGA UCEC and TCGA TARGET GTEx, https://xenabrowser.net/), the University of Alabama at Birmingham (CPTAC, https://ualcan.path.uab.edu/), GDC data portal (TCGA, https://portal.gdc.cancer.gov/), and the Human Protein Atlas (https://www.proteinatlas.org/).

## Statistics

All the statistical tests are described in the figure legends. For Figures 1c, 2j, 2l, and 7k, statistical analysis was done using rstatix or ggpubr packages in R. For supplementary figures 5c, 5d, 5f, 5g, statistical analysis was performed using ggpubr package in R. In Figure 3d, statistical test was conducted using DESeq2 package in R. In Figure 2c, survminer package in R was used for statistical test. For supplementary Figures 1g, 1j, and Figure 7l, 7m, 7o, p values were extracted from UCEC Xena Browser. For supplementary Figures 6a, 6b, p values were extracted from Human Protein Atlas. For all other figures, statistical analysis was conducted using GraphPad Prism software. Results were considered statistically significant if *P* <0.05.

## Acknowledgements

This project was supported by NIH R37CA238274 (SY), U01CA272424 (BL, SY) Administrative Supplements Award, the Department of Pathology Start-Up Fund (SY), the Holden Comprehensive Cancer Center at The University of Iowa and its National Cancer Institute Award P30CA086862.

## Author Contributions

S.Y. administrated the project. K.S., J.H., B.L., and S.Y. designed the study. K.S., and J.H. performed the experiments and data analysis. R.J., W.M., T.L., E.J., E.Z., A.A., J.G., R.M., E.S., C.O., C.G., D.M., M.Y., H.K., X.M., Y.X. performed the experiments and provide technical support. K.S., W.M., S.Y., and R.J. wrote the manuscript. K.S., and S.Y. organized and supervised the study.

