## Supplementary Materials for "Oncogene SETDB1’s Dual Role in Endometrial Cancer: Driving Tumor Progression and Immune Escape"

1. Supplementary Fig. 1:  Supplementary Fig 1: Elevated SETDB1 Expression Across 19 Tumor Types: Highest Levels in EC Among 8 Tumor Types, Correlating with Poorer Survival Outcomes.
2. Supplementary Figure 2: SETDB1 depletion demonstrates slowed cell proliferation in various knockout single clones.
3. Supplementary Fig 3: SETDB1 depletion in 14 clones across six EC cell lines decreases cancer cells growth in vitro.
4. Supplementary Fig. 4: SETDB1 depletion shows reduced tumor growth in Ishikawa and ECC1 prior injections.
5. Supplementary Fig 5: SETDB1 Target Genes in More Clones, and Differential Expression Analysis of Group 1 and Group 2 Genes Using Two EC Datasets.
6. Supplemental Fig. 6: Prognosis Outcomes of Group 1 and Group 2 genes.
7. Supplementary Figure 7: Loss of H3K9me3 Binding on Many ZNFs, but Not on SETDB1 Itself in SETDB1-/- Cells, Evidenced by ChIP-PCR Analysis of H3K9me3 on ZNF266, ZNF841, and ZNF582 Promoters.
8. Supplementary Figure 8: Knockout SETDB1 leads to defect in chromosome segregation.
9. Supplementary Fig. 9: SETDB1 regulates repeat elements and interferon pathways activation in endometrial cancer.
10. Supplementary Fig 10. CCL5 and CXCL10 is the most upregulated cytokines in SETDB1-/- tumors.

**Supplementary Fig 1: Elevated SETDB1 Expression Across 20 Tumor Types: Highest Levels in EC Among 8 Tumor Types, Correlating with Poorer Survival Outcomes**

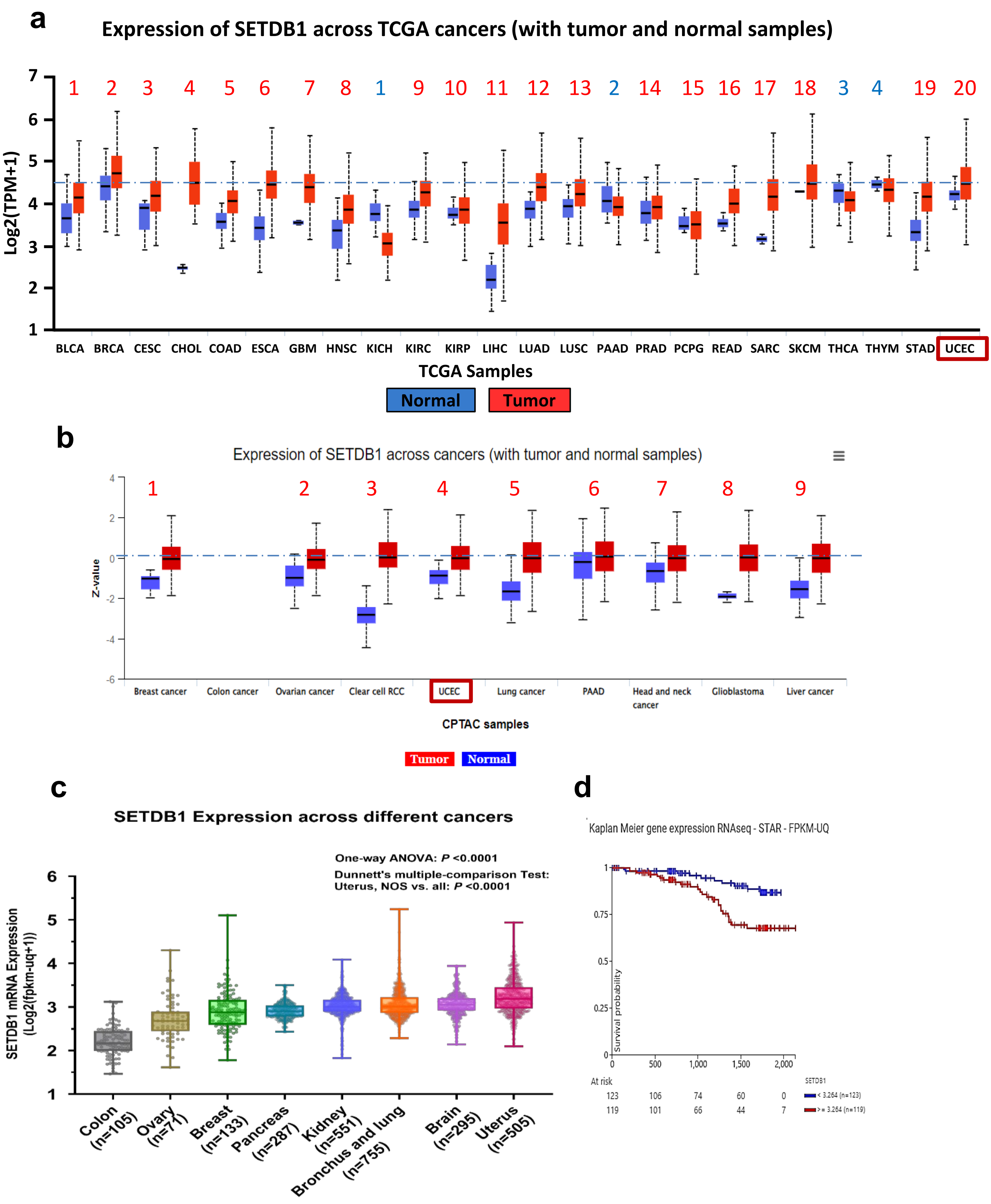

**Supplementary Figure 2: SETDB1 Depletion in 17 Clones Across Three Endometrial Cancer Cell Lines Demonstrates Slowed Cell Proliferation in Initial Screening.**

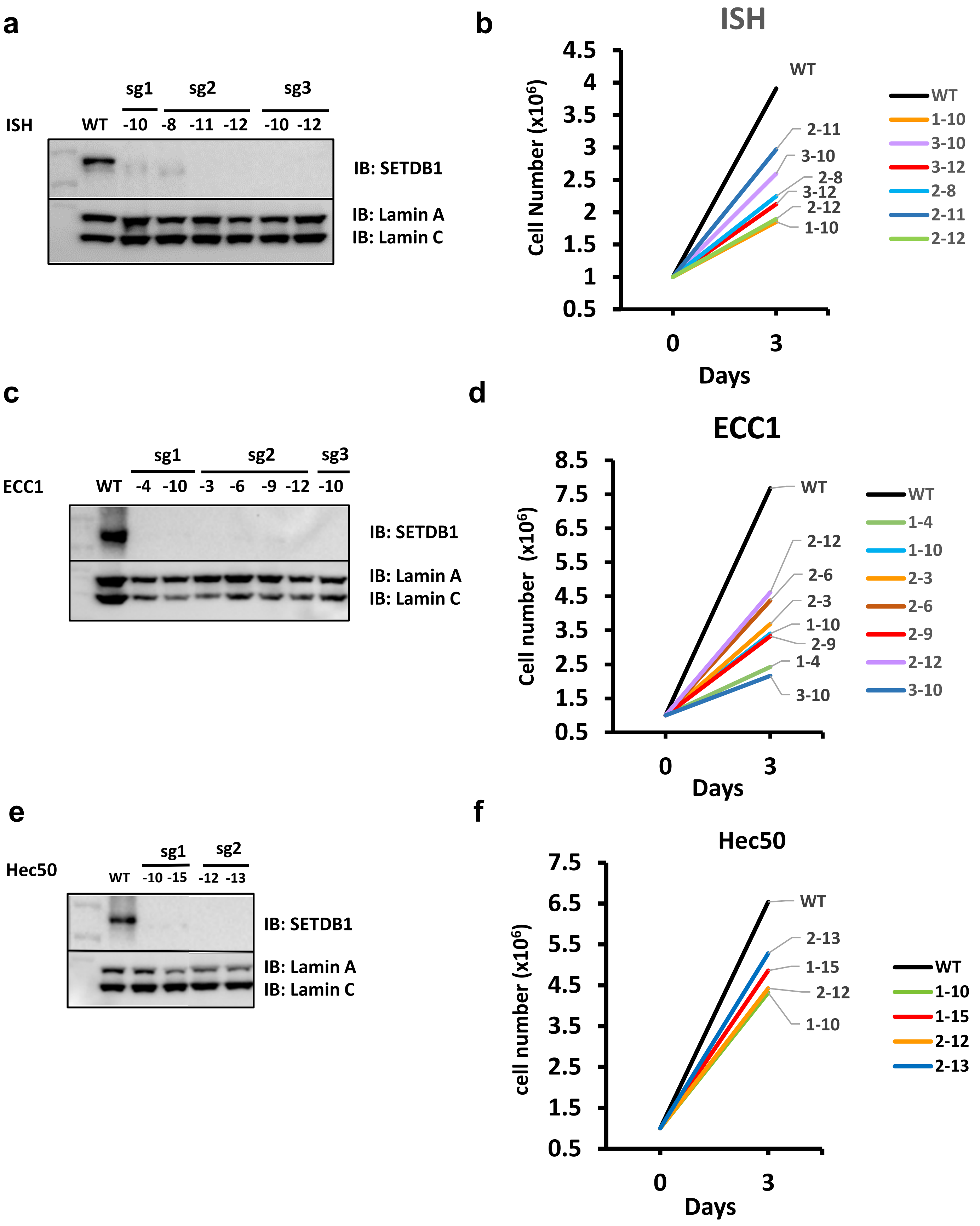

**Supplementary Fig 3: SETDB1 depletion in 14 clones across six EC cell lines decreases cancer cells growth in vitro.**

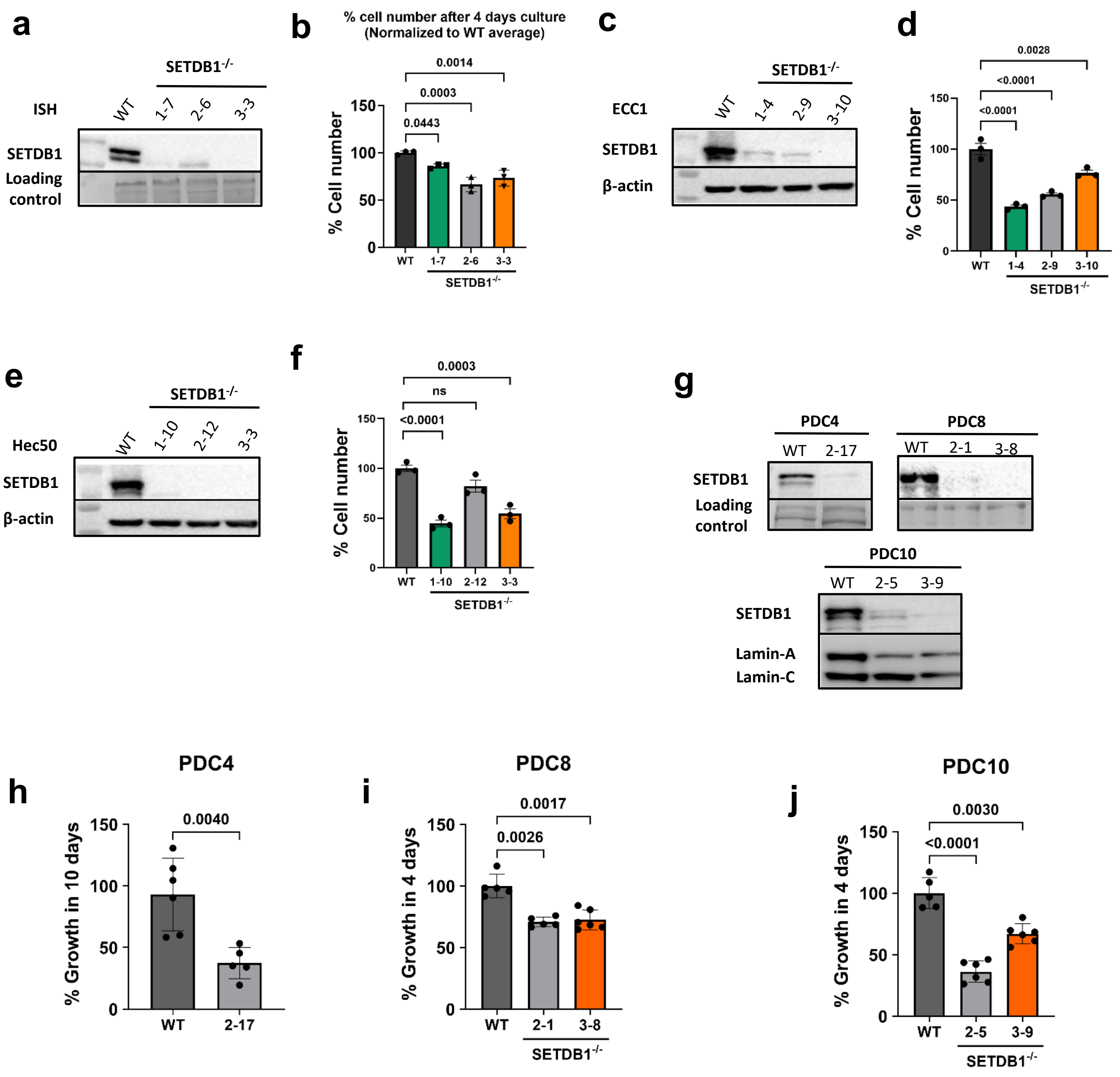

**Supplementary Fig. 4: SETDB1 depletion shows reduced tumor growth in Ishikawa and ECC1 prior injections**

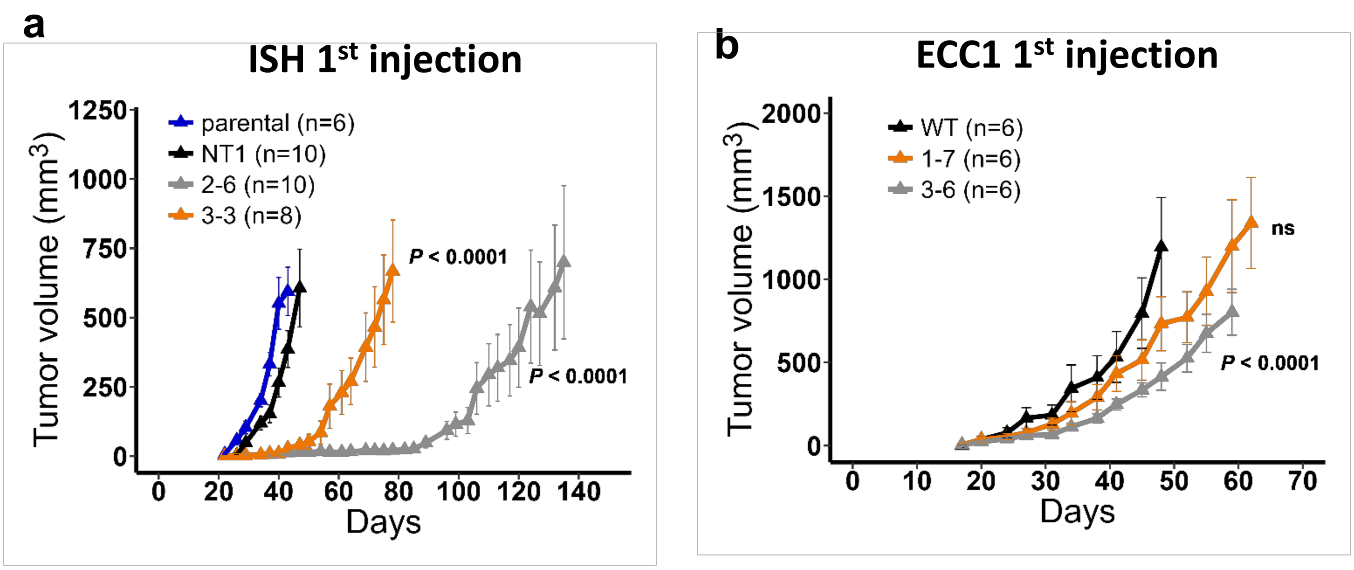

**Supplementary Figure 5: SETDB1 Target Genes in More Clones, and Differential Expression Analysis of Group 1 and Group 2 Genes Using Two EC Datasets.**

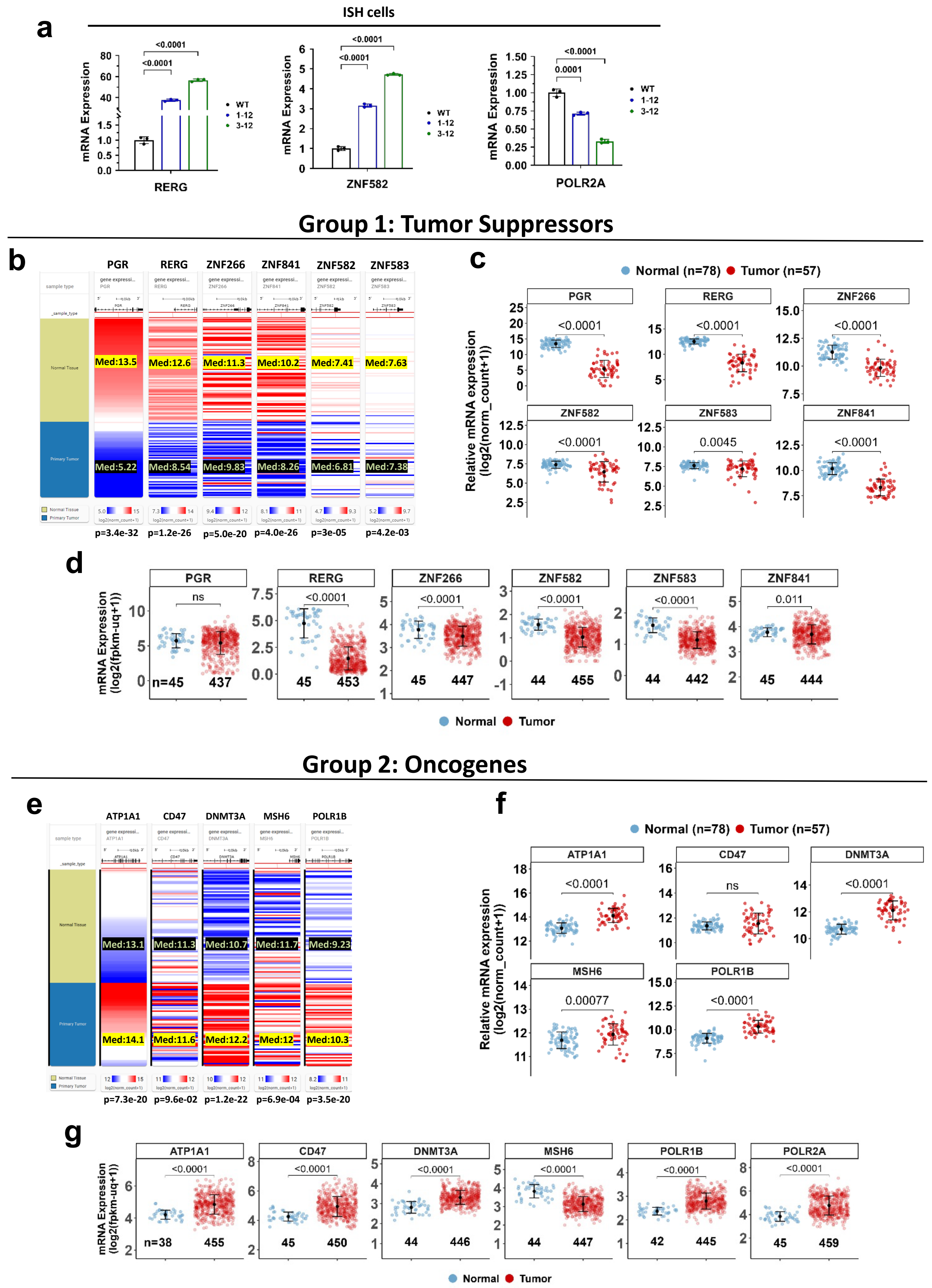

**Supplemental Figure 6: Prognosis Outcomes of Group 1 and Group 2 genes.**

**a** **Group 1 (Tumor suppressors repressed by SETDB1)**

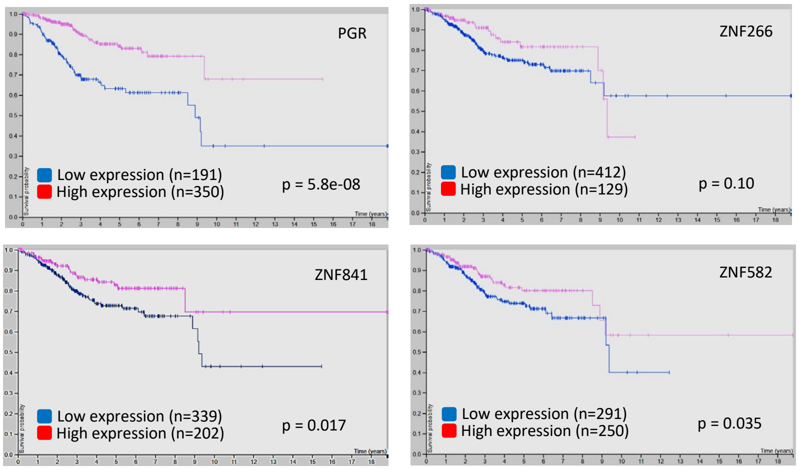


**b** **Group 2 (Oncogenes promoted by SETDB1)**

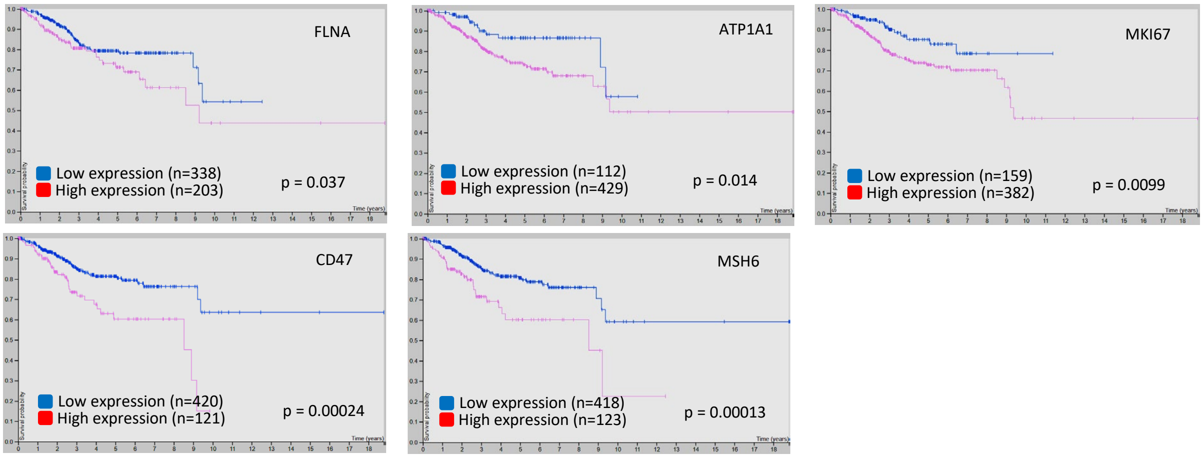

**Supplementary Figure 7: Loss of H3K9me3 Binding on Many ZNFs, but Not on SETDB1 Itself in SETDB1-/- Cells, Evidenced by ChIP-PCR Analysis of H3K9me3 on ZNF266, ZNF841, and ZNF582 Promoters**

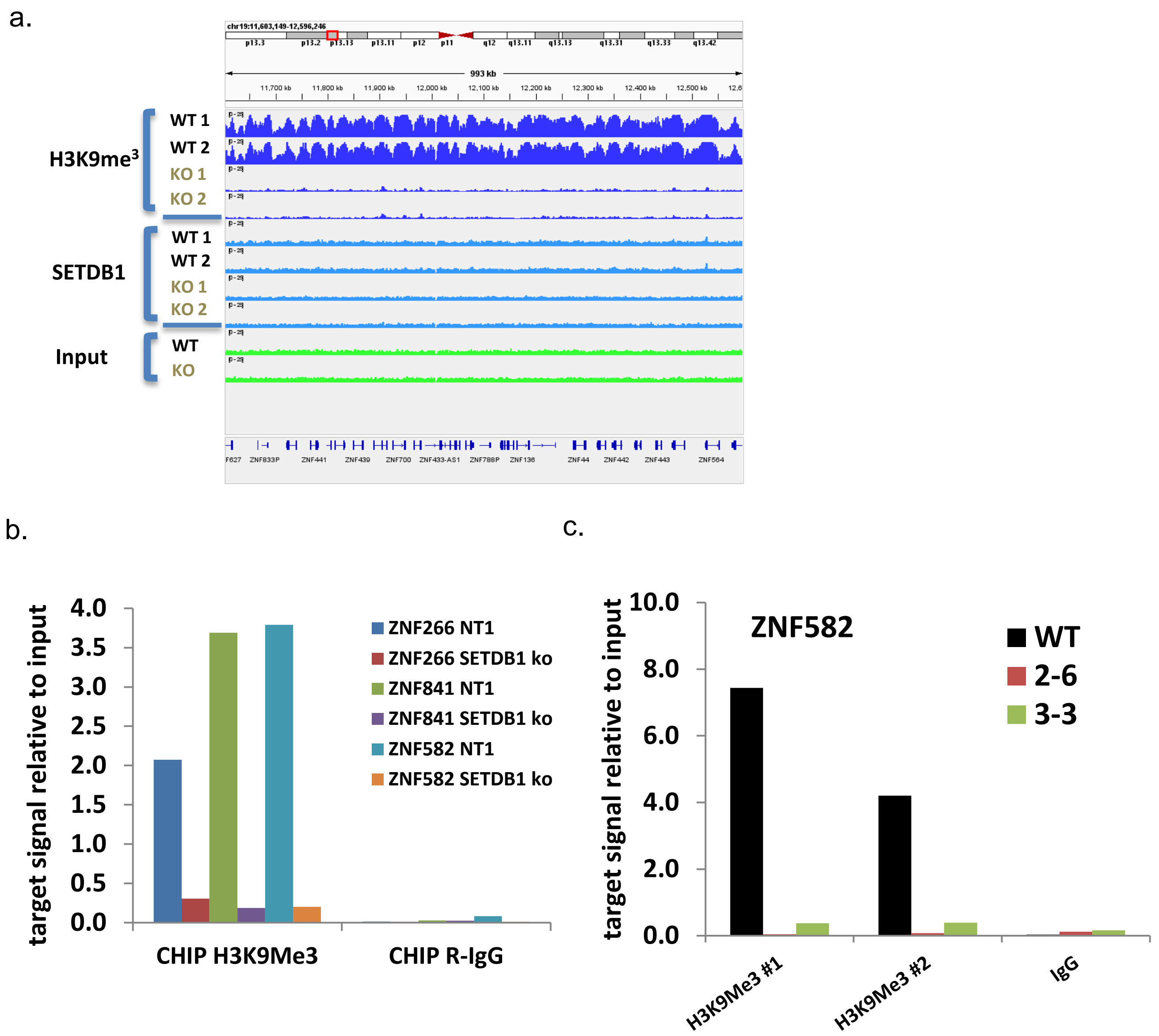

#### Supplementary figure 8. Knockout SETDB1 leads to defect in chromosome segregation

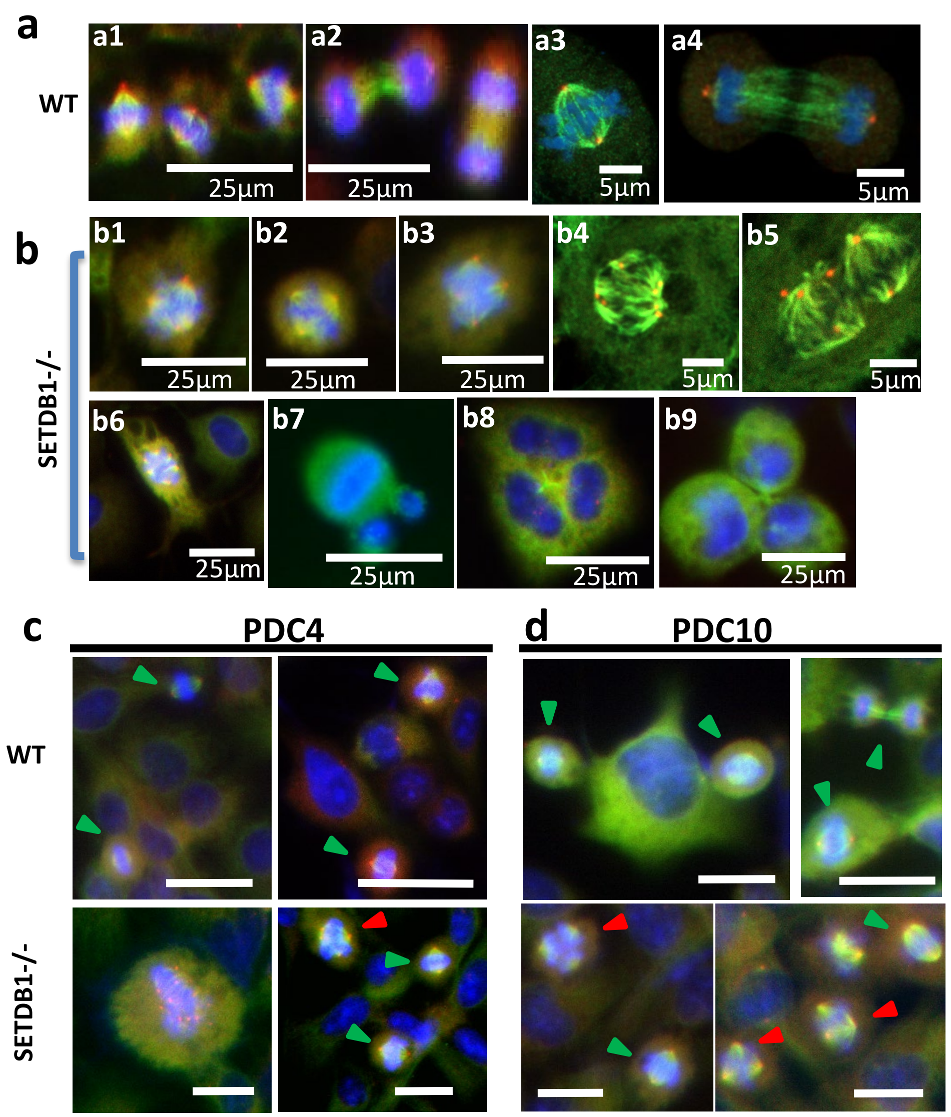

### Supplementary Figure 9: SETDB1 regulates repeat elements and interferon pathways activation in endometrial cancer

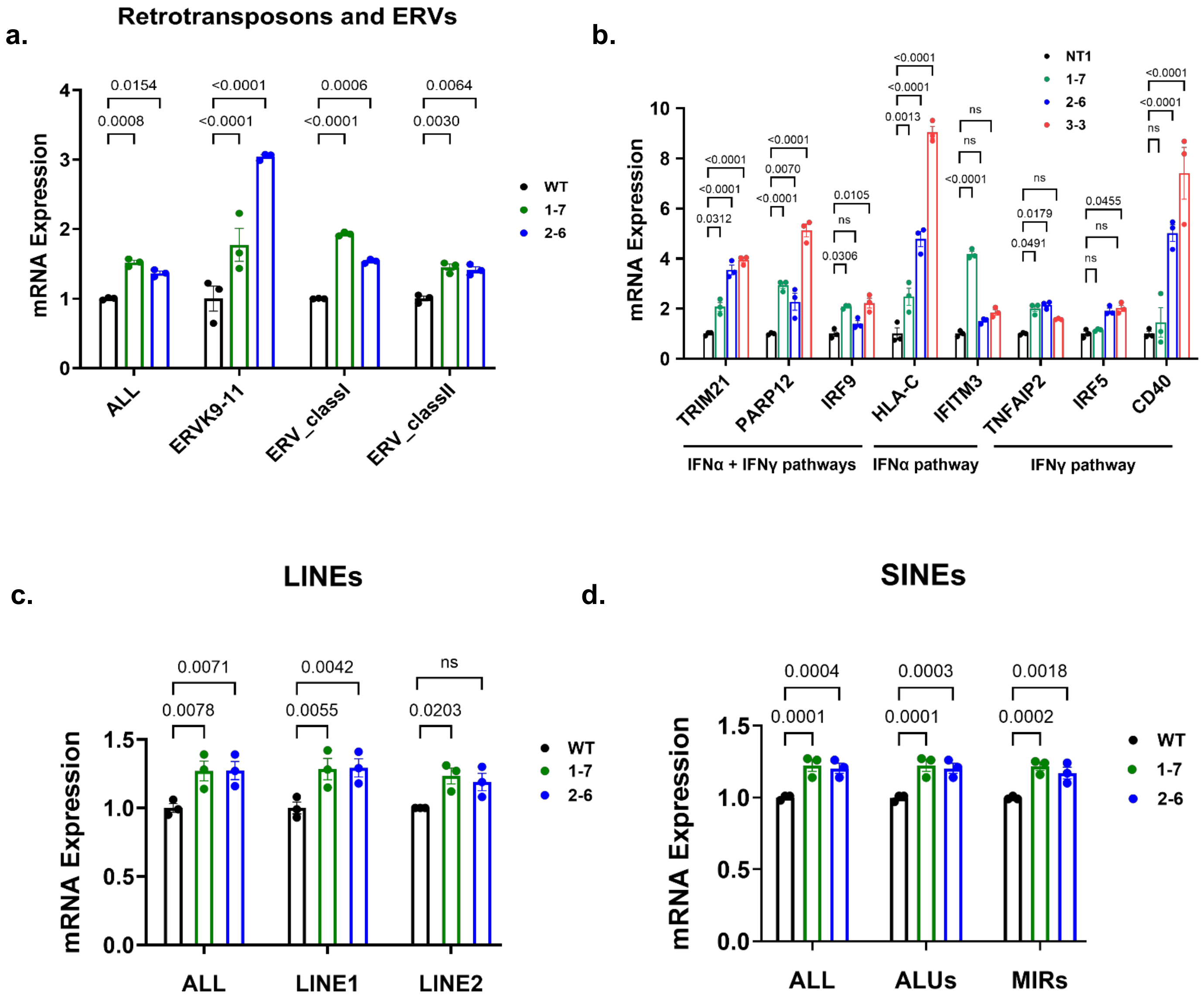

### Supplementary Figure 10. CCL5 and CXCL10 are the most upregulated cytokines in SETDB1$^{-/-}$ tumors.

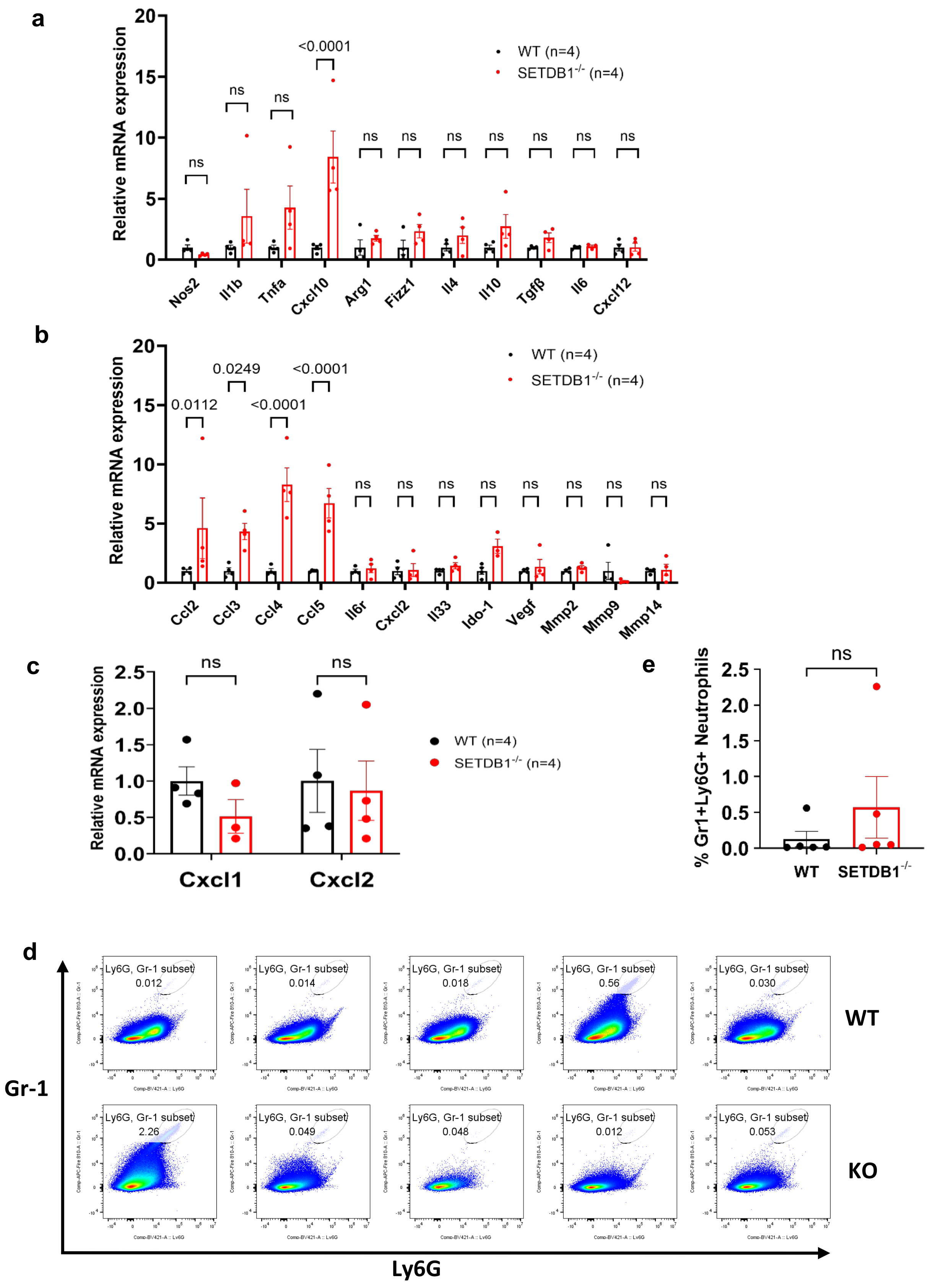

Supplementary materials

Supplementary Table 1.

Human primers:

| Primer name | Primer Sequence |
|---|---|
| PGR-For | ACTGGGTTTGACTTCGTAGCCCTT |
| PGR-Rev | ATGTGGCAGATCCCACAGGAGTTT |
| RERG-For | TGGTCTACGACATTACTGACCG |
| RERG-Rev | AAGCACAAGCCAATTCTGTGG |
| ZNF266-For | AAACCAAAGAGTTAGCCCTTCAG |
| ZNF266-Rev | GGCTTCCTATCATTTGAATCCCA |
| ZNF841-For | TTCAGTCCGGTCTGGGTGAAT |
| ZNF841-Rev | CTCGTCTTGTGTAGGTAACGAAA |
| ZNF582-For | ATGTCCCTTGGGTCAGAATTGT |
| ZNF582-Rev | TTGCCTTGCTCTAGGAAGGAG |
| FLNA-For | GGAGGAGGCAAAAGTGACCG |
| FLNA-Rev | ACTTATCCACGTACACCTCGAAG |
| FLNB-For | GTGAACAAACGCATCGGCAA |
| FLNB-Rev | ACCAGACCCAAGATGAGCTTC |
| JAG1-For | GCCGAGGTCCTATACGTTGC |
| JAG1-Rev | CCGAGTGAGAAGCCTTTTCAA |
| POLR2A-For | GGGTGGCATCAAATACCCAGA |
| POLR2A-Rev | AGACACAGCGCAAAACTTTCA |
| CD47-For | AGAAGGTGAAACGATCATCGAGC |
| CD47-Rev | CTCATCCATACCACCGGATCT |
| MSH6-For | CCAAGGCGAAGAACCTCAAC |
| MSH6-Rev | ACCAGGGGTAACCCTCCATC |
| MUC5AC-For | ACCAATGCTCTGTATCCTTCCC |
| MUC5AC-Rev | TGGTGGACGGACAGTCACT |
| ATP1A1-For | ACAGACTTGAGCCGGGGATTA |
| ATP1A1-Rev | TCCATTCAGGAGTAGTGGGAG |
| POLR1B-For | TCCGAATGTTGATTATGCCTCG |
| POLR1B-Rev | TGACAGCGGAATGTTCTTCCC |
| CCL4-For | GCTTCCTCGCAACTTTGTGG |
| CCL4-Rev | TCACTGGGATCAGCACAGAC |
| IFNa-For | GCCTCGCCCTTTGCTTTACT |
| IFNa-Rev | CTGTGGGTCTCAGGGAGATCA |
| CCL5-For | CCAGCAGTCGTCTTTGTCAC |
| CCL5-Rev | CTCTGGGTTGGCACACACTT |
| CXCL9-For | CAAGGGACTACTCACCTACAATC |
| CXCL9-Rev | ACATCTGCTGAATCTGGGTTTA |
| IFNg-For | TTGGAAAGAGGAGAGTGACAG |
| IFNg-Rev | ACATTCATGTCTTCCTTGATGG |
| IFNβ-For | TTGACATCCCTGAGGAGATTAAGC |
| IFNβ-Rev | TTAGCCAGGAGGTTCTCAACAATAG |

| | |
|---|---|
| TNFRSF10B-For | GCCCCACAACAAAAGAGGTC |
| TNFRSF10B-Rev | AGGTCATTCCAGTGAGTGCTA |
| GAPDH-For | TCAAGGCTGAGAACGGGAAG |
| GAPDH-Rev | CGCCCCACTTGATTTTGGAG |

Mouse primers:

| **Primer name** | **Primer Sequence** |
|---|---|
| Nos2-For | CACTTGGATCAGGAACCTGAAGCCC |
| Nos2-Rev | CTTTGTGCTGGGAGTCATGGAGCCG |
| Il1β-For | TGGACCTTCCAGGATGAGGACA |
| Il1β-Rev | GTTCATCTCGGAGCCTGTAGTG |
| Tnfα-For | CTGTAGCCCACGTCGTAGC |
| Tnfα-Rev | TTGAGATCCATGCCGTTG |
| Cxcl10-For | CCAAGTGCTGCCGTCATTTTC |
| Cxcl10-Rev | GGCTCGCAGGGATGATTTCAA |
| Arg1-For | AGAGATTATCGGAGCGCCTT |
| Arg1-Rev | TTTTTCCAGCAGACCAGCTT |
| Fizz1-For | CCAATCCAGCTAACTATCCCTCC |
| Fizz1-Rev | ACCCAGTAGCAGTCATCCCA |
| Il4-For | GGTCTCAACCCCCAGCTAGT |
| Il4-Rev | GCCGATGATCTCTCTCAAGTGAT |
| Il10-For | ATCGATTTCTCCCCTGTGAA |
| Il10-Rev | TGTCAAATTCATTCATGGCCT |
| Tgfβ-For | GGAGAGCCCTGGATACCAAC |
| Tgfβ-Rev | CAACCCAGGTCCTTCCTAAA |
| Il6r-For | GGCTCACAAAACAGAGAATGG |
| Il6r-Rev | GGTGATCATTCAGGGAGCAT |
| Cxcl12-For | ATTCTCAACACTCCAAACTGTGC |
| Cxcl12-Rev | ACTTTAGCTTCGGGTCAATGC |
| Ccl2-For | CCCAATGAGTAGGCTGGAGA |
| Ccl2-Rev | TCTGGACCCATTCCTTCTTG |
| Ccl3-For | TTCTCTGTACCATGACACTCTGC |
| Ccl3-Rev | CGTGGAATCTTCCGGCTGTAG |
| Ccl4-For | TTCCTGCTGTTTCTCTTACACCT |
| Ccl4-Rev | CTGTCTGCCTCTTTTGGTCAG |
| Ccl5-For | GCTGCTTTGCCTACCTCTCC |
| Ccl5-Rev | TCGAGTGACAAACACGACTGC |
| Cxcl2-For | CCAACCACCAGGCTACAGG |
| Cxcl2-Rev | GCGTCACACTCAAGCTCTG |
| Il33-For | TCCAACTCCAAGATTTCCCCG |
| Il33-Rev | CATGCAGTAGACATGGCAGAA |
| Ido-1-For | TGGCGTATGTGTGGAACCG |
| Ido-1-Rev | CTCGCAGTAGGGAACAGCAA |
| Vegf-For | GCTTCCTACAGCACAGCAGA |
| Vegf-Rev | AATGCTTTCTCCGCTCTGAA |
| Mmp2-For | TCTGCGATGAGCTTAGGGAAAC |
| Mmp2-Rev | GACATACATCTTTGCAGGAGACAAG |

| | |
|---|---|
| Mmp9-For | ACGACATAGACGGCATCCAGTATC |
| Mmp9-Rev | AGGTATAGTGGGACACATAGTGGG |
| Mmp14-For | AGGCTGATTTGGCAACCATG |
| Mmp14-Rev | CCCACCTTAGGGGTGTAATTCTG |
| Cxcl1-For | CTGGGATTCACCTCAAGAACATC |
| Cxcl1-Rev | CAGGGTCAAGGCAAGCCTC |
| Gapdh-For | CTTCAACAGCAACTCCCACTCTTCC |
| Gapdh-Rev | GGTGGTCCAGGGTTTCTTACTCC |

ChIP primers:

| | |
|---|---|
| ZNF582-For | GCGAGATCCGGCTTCAA |
| ZNF582-Rev | CAGACGCACAAAGCACAC |
| ZNF841-For | ACGTCGCTATGAGTGTGTTTC |
| ZNF841-Rev | GGGCTGTTGCGGGTATTT |
| ZNF266-For | CCTGGGTAACATACCTCAACTC |
| ZNF266-Rev | GGCCTATCTCCTCCATTCATTC |

Supplementary Figure Legends:

Supplementary Figure 1. **Elevated SETDB1 Expression Across 20 Tumor Types: Highest Levels in EC Among 8 Tumor Types, Correlating with Poorer Survival Outcomes. a.** TCGA SETDB1 mRNA Expression comparison between normal (n=) and tumor (n=) tissues across 20 different cancer types. **b.** Expression of SETDB1 protein in normal and tumor tissues across 9 different cancers in CPTAC dataset. Colon cancer had no available data. **c.** TCGA SETDB1 mRNA expression across 8 different tumor tissues: Colon (n=105), Ovary (n=71), Breast (n=133), Pancreas (n=287), Kidney (n=551), Bronchus and lung (n=755), Brain (n=295), and Uterus (n=505). **d.** Kaplan Meier survival curve from TCGA dataset illustrating prognostic outcomes for SETDB1 high (n=119) and SETDB1 low (n=123) patients' group. In **a, b, c**, boxplots are expanded from min to max values of the datapoints. The low and high end of the box represent 25th and 75th percentiles respectively. Middle horizontal line represents the median. Statistical tests: One-way ANOVA test followed by Dunnet's multiple comparison test (**c**) Log-Rank test (**d**).

Supplementary Figure 2. **SETDB1 Depletion in 17 Clones Across Three Endometrial Cancer Cell Lines Demonstrates Slowed Cell Proliferation in Initial Screening. a, c, e.** Western blot analysis for SETDB1 protein conducted on SETDB1 knockout single clones generated for Ishikawa (n=6), ECC1 (n=7), Hec50 (n=4) respectively compared with their wildtype (WT) counterparts. **b, d, f.** cell proliferation rates for individually generated knockout single clones from Ishikawa, ECC1, and Hec50 cell lines respectively.

Supplementary Figure 3. **SETDB1 depletion in 14 clones across six EC cell lines decreases cancer cells growth in vitro. a, c, e.** SETDB1 immunoblots for wildtype and SETDB1 knockout single clones in Ishikawa, ECC1 and Hec50 cell lines respectively. **b, d, f.** % normalized cell counts to wildtype for Ishikawa, ECC1, and Hec50 SETDB1 knockouts respectively 4 days after plating in culture dish (n=3 biological replicates). **g.** Western blots for generated SETDB1-/- PDC cells (PDC4, 8 and 10). Loading control represents Coomassie blue stained gels. **h.** % Growth of PDC4 wildtype and SETDB1-/- clone in 10 days calculated through normalized absorbance readings from resazurin assay (n=5-6 biological replicates). **i, j.** % Growth of PDC8 and PDC10 wildtype and SETDB1-/- clone respectively in 4 days calculated through normalized absorbance readings from resazurin assay (n=5-6 biological replicates). **b,d,f, h-j**, Data shown as mean ± SD. P values are calculated using one-way ANOVA followed by post hoc Dunnett's multiple comparison test (**b,d,f**) Welch's t test (**h**) and one-way ANOVA test followed by Post hoc Dunnett's T3 multiple comparisons test (**i, j**).

Supplementary Figure 4. **SETDB1 depletion shows reduced tumor growth in Ishikawa and ECC1 prior injections. a, b.** Mean tumor volume measurements over time from first injection experiment on female NSG mice for wildtype and SETDB1 knockout clones belonging to cancer cells: Ishikawa (n=6/group for WT, n=10/group for NT1, 2-6, and n=8/group for 3-3) (**a**), ECC1 (n=6/group for WT, 1-7, 3-6) (**b**) with n representing number of total tumors. Data shown as mean ± SEM. P values were calculated through a non-linear fit (Exponential growth model as log(population)) to the curves and the fitted curves were compared using extra sum-of-squares F test.

Supplementary Figure 5. **SETDB1 Target Genes in More Clones, and Differential Expression Analysis of Group 1 and Group 2 Genes Using Two EC Datasets. a.** Verified qPCR results for Group 1 and Group 2 genes mRNA expression on additional Ishikawa SETDB1-/- clones (n=3 technical replicates). **b, e.** Heatmaps illustrating mRNA expression comparison across normal and tumor tissues in EC-TCGA TARGET GTEx dataset for Group 1 and Group 2 genes respectively (n=57 for Primary Tumor, n=78 for Normal Tissue). **c, f.** Jitter plots illustrating the distribution of mRNA expression across normal and tumor tissues for Group 1 and 2 genes respectively in EC-TCGA TARGET GTEx dataset (n=57 for Primary Tumor, n=78 for Normal Tissue). **d, g.** Jitter plots illustrating the distribution of mRNA expression across normal and tumor tissues for Group 1 and 2 genes respectively in EC CPTAC dataset. Value n represents number of patients in each group. Outliers, defined as values falling outside of 1.5 times the Interquartile Range of the first and third quartiles, were omitted from the data points. For **a**, Data shown as mean ± SD. Statistical tests: Student's t-test (**a**), Student's t-test with Bonferroni correction (**c, d, f, g**). ns, not significant.

Supplementary Fig. 6. **Prognosis Outcomes of Group 1 and Group 2 genes. a, b.** Kaplan Meier survival curves for Group 1 and 2 genes respectively. P values were calculated using Log-Rank test.

Supplementary Figure 7: **Loss of H3K9me3 Binding on Many ZNFs, but Not on SETDB1 Itself in SETDB1$^{-/-}$ Cells, Evidenced by ChIP-PCR Analysis of H3K9me3 on ZNF266, ZNF841, and ZNF582 Promoters. a.** ChIP-seq H3K9me3 and SETDB1 peaks on ZNF gene clusters on chromosome 19. **b.** qPCR analysis of H3K9me3 and IgG ChIP samples for ZNF266, 841, and 582 promoter sites on wildtype and SETDB1 knockout cells. **c.** qPCR analysis of H3K9me3 and IgG ChIP samples for ZNF582 promoter site on 2-6 and 3-3 knockout clones and wildtype cells.

Supplementary Figure 8. **Knockout SETDB1 leads to defect in chromosome segregation. a, b** Representative Immunofluorescence staining images of bipolar and multipolar mitotic cells in wildtype and SETDB1 knockout Ishikawa cells respectively. Red: Y-tubulin, green: β-tubulin, blue: genomic DNA. **c, d** Representative Immunofluorescence staining images of bipolar and multipolar mitotic cells in wildtype and SETDB1 knockout PDC4, 10 cell lines respectively. Red: Y-tubulin, green: β-tubulin, blue: genomic DNA. Scale bars represent 25um.

Supplementary Figure 9. **SETDB1 regulates repeat elements and interferon pathways activation in endometrial cancer. a.** mRNA expression across different classes of Retrotransposons and ERVs and their overall (ALL) expressions in SETDB1$^{-/-}$ and wildtype Ishikawa cells (n=3 biological replicates). **b.** mRNA expression analysis of genes related to IFNα and IFNγ pathways in SETDB1$^{-/-}$ and wildtype Ishikawa cell line (n=3 biological replicates). **c, d.** Quantification of mRNA expression for different classes of LINEs and SINEs repeat elements respectively and their overall (ALL) expressions in SETDB1$^{-/-}$ and wildtype Ishikawa cells (n=3 biological replicates). Data shown as mean ± SD. P values were calculated using Two-way ANOVA followed by post hoc Dunnett's multiple comparison test. ns, not significant.

Supplementary Figure 10. **CCL5 and CXCL10 is the most upregulated cytokines in SETDB1$^{-/-}$ tumors. a, b.** mRNA expression quantification across several murine cytokines and markers in SETDB1$^{-/-}$ and wildtype Ishikawa tumors (n=4 per group). **c.** Quantification of neutrophils recruiting Cxcl1 and Cxcl2 mice cytokines mRNA expression in SETDB1$^{-/-}$ and wildtype Ishikawa tumors (n=4 per group). **d.** Representative gating strategy for Gr1+Ly6G+ neutrophils in scatter plot for SETDB1$^{-/-}$ and wildtype Ishikawa tumors (n=5 per group). **e.** Percent Gr1+Ly6G+ neutrophils (out of total CD45+ immune cells) in SETDB1$^{-/-}$ and wildtype Ishikawa tumors (n=5 per group). For **a-c** and **e**, Data shown as mean ± SEM. Statistical tests: Two-way ANOVA with post hoc Sidak multiple comparison test (**a-c**) Unpaired Welsh's t-test (**e**). ns, not significant.
